# Unsupervised cell functional annotation for single-cell RNA-Seq

**DOI:** 10.1101/2021.11.20.469410

**Authors:** Dongshunyi Li, Jun Ding, Ziv Bar-Joseph

## Abstract

One of the first steps in the analysis of single cell RNA-Sequencing data (scRNA-Seq) is the assignment of cell types. While a number of supervised methods have been developed for this, in most cases such assignment is performed by first clustering cells in low-dimensional space and then assigning cell types to different clusters. To overcome noise and to improve cell type assignments we developed UNIFAN, a neural network method that simultaneously clusters and annotates cells using known gene sets. UNIFAN combines both, low-dimensional representation for all genes and cell specific gene set activity scores to determine the clustering. We applied UNIFAN to human and mouse scRNA-Seq datasets from several different organs. As we show, by using knowledge on gene sets, UNIFAN greatly outperforms prior methods developed for clustering scRNA-Seq data. The gene sets assigned by UNIFAN to different clusters provide strong evidence for the cell type that is represented by this cluster making annotations easier.

**Software:** https://github.com/doraadong/UNIFAN

## Introduction

The large increase in studies profiling RNA-Sequencing data in single cells (Tanay and Regev 2017) raises several computational challenges. One of the first, and most important, steps in the analysis of such studies is cell type assignment (Clarke et al. 2021). Several methods have been developed for such assignment, including (semi-)supervised and unsupervised methods. (Semi-)supervised methods mainly use previously annotated datasets to annotate new datasets (Abdelaal et al. 2019). This is done by directly classifying each cell (Alavi et al. 2018; Pliner et al. 2019), by learning an alignment between the datasets (Kiselev et al. 2018) or by joint training using multiple datasets to classify groups of cells in a new study (Brbić et al. 2020; Hu et al. 2020).

While supervised methods are useful in some cases, they cannot be applied to all cases since reference datasets are not available for most organs, tissues and conditions. Another challenge with supervised methods is their inability to identify new cell types, which is often one of the major goals of the study (Abdelaal et al. 2019). Thus, the most popular way to annotate single cell data is by using unsupervised methods. These are often based on clustering cells in a low-dimensional space and manually annotating each cluster using known marker genes or cluster specific differentially expressed genes. Several methods for clustering single cell data have been developed and used. These include SIMLR (Wang et al. 2017) which clusters cells by using multiple kernel functions to construct a similarity matrix between cells, Leiden clustering (Traag et al. 2019) and Seuratv3 (Stuart et al. 2019) which use k-nearest neighbors (k-nn) based graph partitions to group cells, SCCAF (Miao et al. 2020) which refines initial clusters using a self-projection-based approach, and methods based on deep neural networks, such as DESC (Li et al. 2020), which uses autoencoders to reduce the dimensions of the data and then clusters cells in the reduced dimension space.

While several clustering methods have been developed and used for scRNA-Seq data, to date these methods have only relied on the observed expression data. However, there are several additional complementary datasets that can be used to improve clustering and reduce noise related grouping. Specifically, gene sets (Subramanian et al. 2005) have been compiled to characterize many processes, pathways and conditions. While the exact processes or functions that are activated in specific cells or clusters are unknown, we can use these sets to guide the grouping of cells by placing more emphasis on co-expression of genes in known sets when clustering single cell data. Since cells of the same type likely share many of the biological processes, such design can both, improve the identification of good clusters and help in annotating them based on the function of the sets associated with each cluster.

Here we introduce UNIFAN (**Un**supervised S**i**ngle-cell **F**unctional **An**notation) to simultaneously cluster and annotate cells with known biological processes (including pathways). For each cell, we first infer its gene set activity scores based on the co-expression of genes in known gene sets. We also use an autoencoder that outputs a low-dimensional representation learned from the expression of all genes. We combine both, the low dimension representation and the gene set activity scores to determine the cluster for each cell. The process is iterative and we define a target function and show how to learn model parameters to optimize it. In addition to the cell clusters, the method also outputs the gene sets associated with each cluster and these can be used to annotate and assign cell types to different clusters.

We applied UNIFAN to several mouse and human datasets spanning multiple organs, cell types and labs. Our results indicate that by using gene sets as input, UNIFAN can improve on current single cell clustering methods. In addition, in most cases, the gene sets selected for each cluster serve as a very good source for their annotations.

## Results

We developed UNIFAN (**Un**supervised S**i**ngle-cell **F**unctional **An**notation) to simultaneously cluster and annotate cells (and cell clusters) with known biological processes or pathways. By integrating prior information about gene sets with observed expression data, UNIFAN can improve clustering results while simultaneously making the clusters more interpretable. Figure 1 presents an overview of UNIFAN. The method starts with inferring gene set activity scores for each single cell, based on the expression levels of genes in known gene sets. Next, UNIFAN clusters cells by using the learned gene set activity scores and a low-dimensional representation of the expression of all genes in the cell. This is done using an autoencoder-based neural network. The “annotator” part of this network uses the gene set activity scores to guide the clustering such that cells sharing similar biological processes are more likely to be grouped together. See Methods for details on the architecture of UNIFAN and on how parameters are learned for UNIFAN.

**Figure 1:**
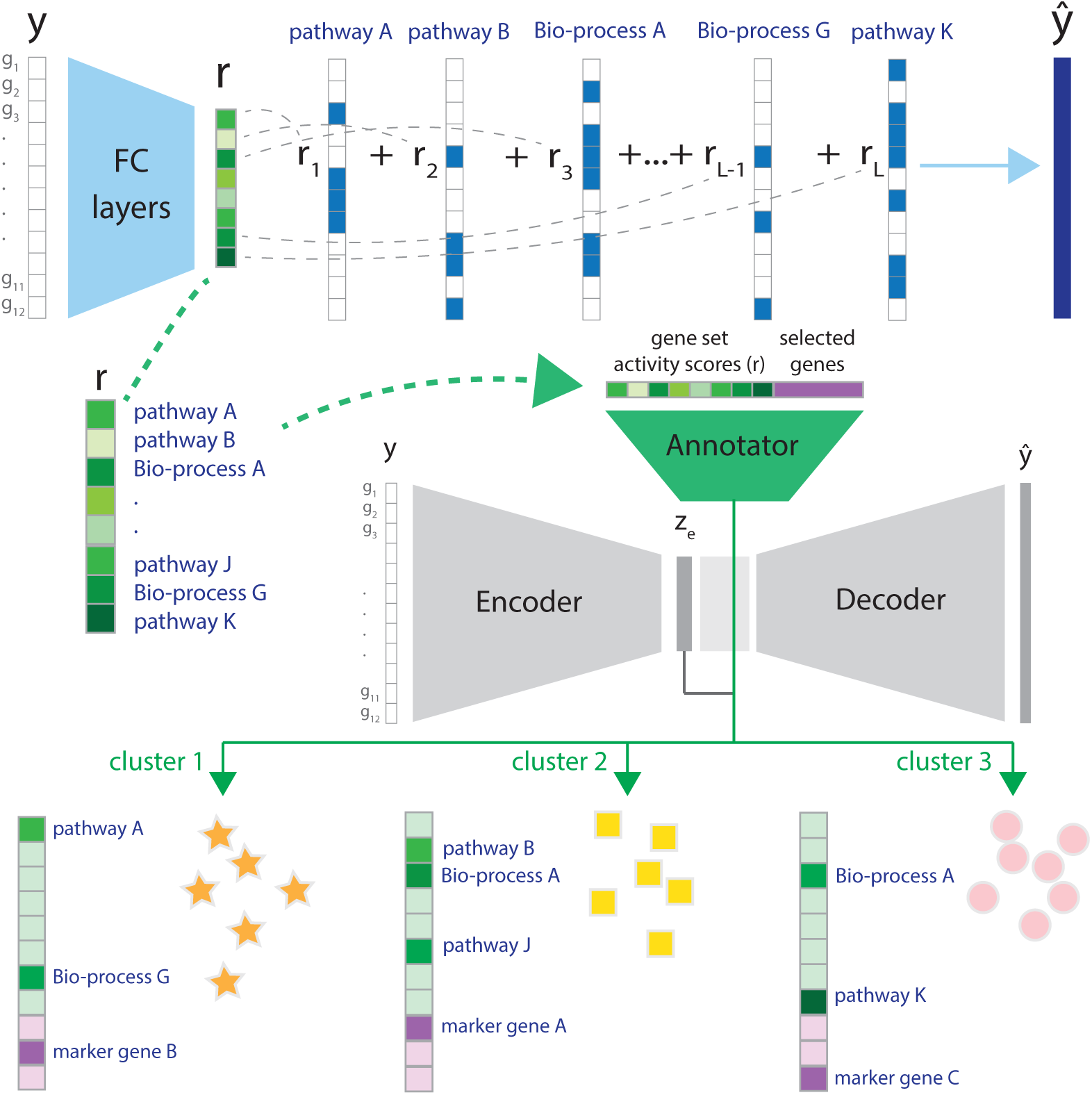
Overview of UNIFAN. **Top:** Using the expression levels for genes in a cell **y**, UNIFAN first infers gene set activity scores (**r**), using an autoencoder. The decoder is composed of binary vectors with values indicating if a gene belongs to a known gene set or not. **Middle:** Next, UNIFAN clusters cells by using the learned gene set activity scores and a low-dimensional representation of the expression of all genes in the cell (**z**_**e**_). For this it uses an autoencoder-based neural network, which contains two parts: the cluster assignment part (grey) and the “annotator” (green). The cluster assignment part assigns a cell to clusters based on the low-dimensional representation (**z**_**e**_) while the “annotator” refines clustering and annotates clusters with biological processes and marker genes. **Bottom:** Cells assigned to different clusters characterized by selected gene sets and marker genes. FC layers: fully-connected layers; Bio-process: biological process.

### UNIFAN Correctly Clusters Cells and Identifies Relevant Biological Processes

We first evaluated if UNIFAN can accurately cluster cells and reveal key pathways and cellular functions activated in cells assigned to different clusters. For this, we used the “pbmc28k” scRNA-seq dataset (Methods). UNIFAN clusters successfully captured different cell types when compared to manual annotations (ARI: 0.81, NMI: 0.77). Figure 2 presents UMAP (Becht et al. 2019) visualizations of **z**_**e**_ output from UNIFAN for each cell. As can be seen, clusters are mostly composed of cells from the same type which is a large improvement over other methods including Leiden clustering (shown in Figure 2C), as we discuss below. By relying on known gene sets, UNIFAN is robust to noise and mainly focuses on relevant co-expressed sets of genes leading to much more coherent clusters. We observed similar results for the other datasets we tested as can be seen in Figures S6 - S8.

**Figure 2:**
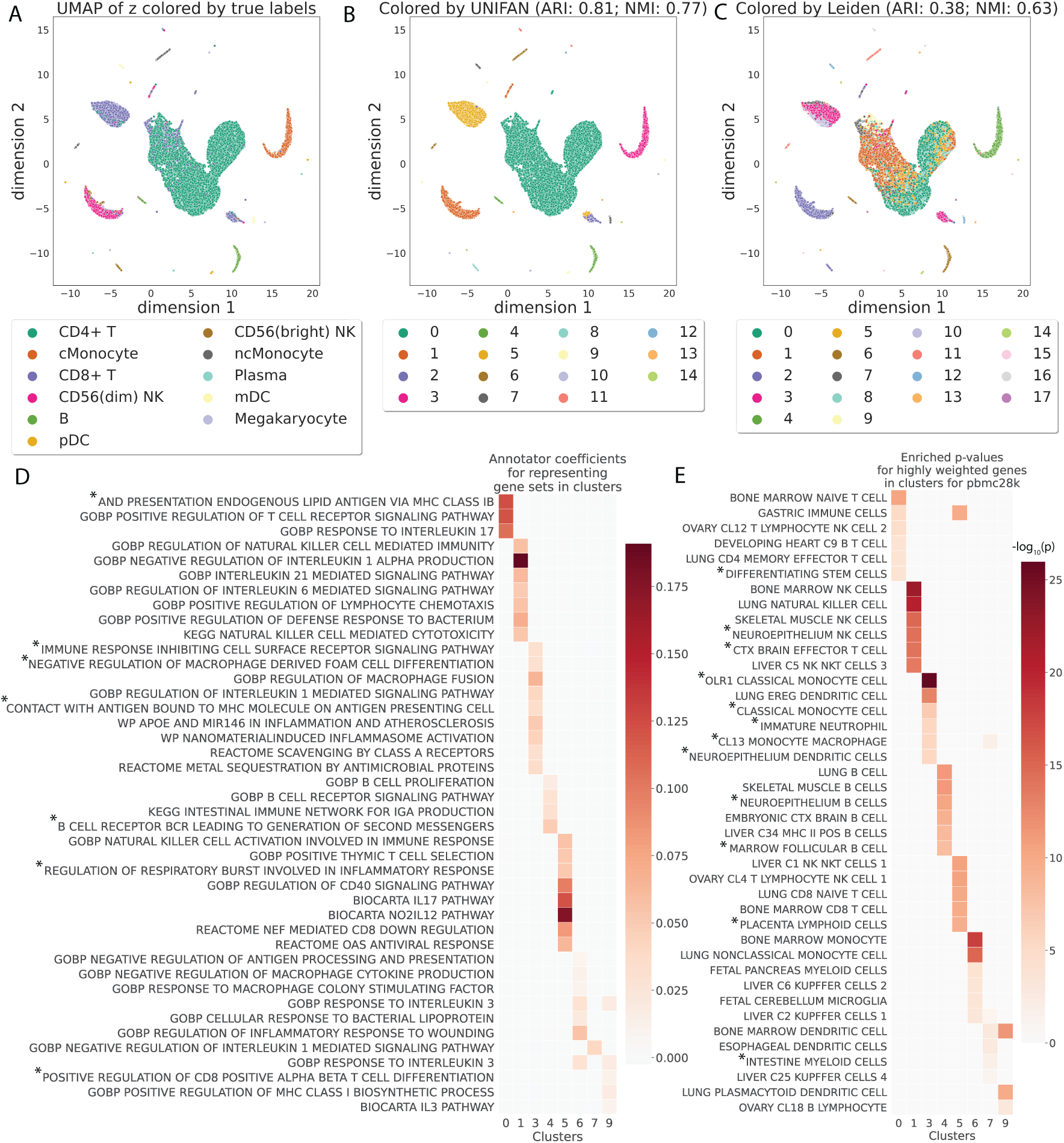
UNIFAN accurately clusters cells and correctly identifies biological processes / pathways. Results presented for the “pbmc28k” dataset. **A, B and C**: UMAP visualization of the low-dimensional representation **z**_**e**_ of cells output from UNIFAN. **A**: Colored by true cell type labels; **B**: colored by the clusters found by UNIFAN; **C**: colored by Leiden clustering. **D**: Coefficients learned by the annotator for highly ranked gene sets for some of the clusters. **E**: Enrichment p-values of cell type marker sets in the highly weighted genes learned by the annotator. Here we show the result from the best run for both UNIFAN and Leiden. Due to space limit, some gene set names in D and E are truncated (marked with *). See Table S2 and S3 for the full names.

To annotate cell clusters, we examined the coefficients assigned by the “annotator” to different gene sets for each cluster. Figure 2D presents some of top ranked sets for the different clusters. We observe that for cluster 0, the set “GOBP POSITIVE REGULATION OF T CELL RECEPTOR SIGNALING PATHWAY” is assigned a large weight and this cluster is annotated as CD4+ T cells in the original paper. For cluster 5 (which mainly contains CD8+ T cells), one of the top scoring sets is “REACTOME NEF MEDIATED CD8 DOWN REGULATION”. Cluster 1 cells labeled as CD56 (dim) natural killer (NK) cells in the original paper. UNIFAN correctly assigns “GOBP REGULATION OF NK CELL MEDIATED IMMUNITY” and “KEGG NK CELL MEDIATED CYTOTOXICITY” as two of the top gene sets for this cluster. Cluster 3 and 6 correspond to classical monocyte (cMonocyte) and non-classical monocyte (ncMonocyte), respectively. While UNIFAN assigns biological processes related to “antigen presentation” and “inflammation” to both clusters, the biological process related to wound healing “GOBP REGULATION OF INFLAMMATORY RESPONSE TO WOUNDING” only appears in cluster 6. One of the main differences between ncMonocyte and cMonocyte is their role in wound healing (Schmidl et al. 2014) and so such assignment can make it much easier to correctly annotate this cluster of cells. In addition to the gene sets, we also evaluated genes highly weighted by the annotator by comparing them to known cell type marker sets. As shown in Figure 2E, the most enriched cell type marker sets for each cluster correspond very well to the true cell labels, indicating that UNIFAN can indeed identify the marker genes for each cell type (cluster).

We observed similar performance in terms of cluster annotations for other datasets we tested. For example, for the “Atlas lung” dataset, UNIFAN successfully separated macrophage (cluster 2) and alveolar macrophage (cluster 5) as shown in Figure S6A. The annotator selected “GOBP MACROPHAGE FUSION”, “WP MACROPHAGE MARKERS” and “GOBP NEGATIVE REGULATION OF RESPONSE TO INTERFERON GAMMA” for both clusters. It also selected “GOBP REGULATION OF COLLAGEN FIBRIL ORGANIZATION” for cluster 8, which agrees well with the labels of cells in that cluster (fibroblasts). It selected “GOBP CILIUM MOVEMENT” for cluster 10, again in agreement with the types of cells in this cluster (ciliated). Similarly, the most enriched cell type marker sets for each cluster, learned from the highly weighted genes, corresponded very well to the true cell type labels (Figure S6E).

### UNIFAN Improves upon Prior Methods

We compared UNIFAN’s clustering performance on all datasets with several prior methods proposed for clustering scRNA-Seq data. The number of cells in the datasets we used to compare the methods ranges from 366 (Aorta in Tabula Muris) to 96,282 (“Atlas lung” dataset) and so they can provide a good representation of the scRNA-Seq datasets being analyzed by researchers. The methods we compared to included two graph-based methods Leiden clustering (Traag et al. 2019) and Seuratv3 (Stuart et al. 2019), a kernel-based method SIMLR (Wang et al. 2017), a self-projection-based method SCCAF (Miao et al. 2020) and a deep-learning based method DESC (Li et al. 2020). In addition to these unsupervised methods, we also compared UNIFAN to two (semi-)supervised deep-learning methods MARS (Brbić et al. 2020) and ItClust (Hu et al. 2020). We allow these two (semi-)supervised methods to use the true labels for 5% of the cells in a dataset for model learning. We also compared to CellAssign (AW Zhang et al. 2019) which uses known cell type markers for cell type assignment. For each dataset, we ran each method ten times using different initializations. Results are presented in Figure 3 and S9. As can be seen, for all datasets, UNIFAN outperforms all other unsupervised methods regardless of the evaluation metric being used (e.g., average ARI/NMI of UNIFAN and the best performing unsupervised prior method on “pbmc28k”: UNIFAN-0.72/0.74, Leiden-0.37/0.62; on “HuBMAP spleen”: UNIFAN-0.75/0.71, DESC-0.31/0.64; on “Tabula Muris”: UNIFAN-0.70/0.75, SIMLR-0.53/0.65). The large improvement may result from the ability of UNIFAN’s to focus on the more relevant sets of co-expressed genes rather than on co-expression that may results from noise or the large number of genes being profiled.

**Figure 3:**
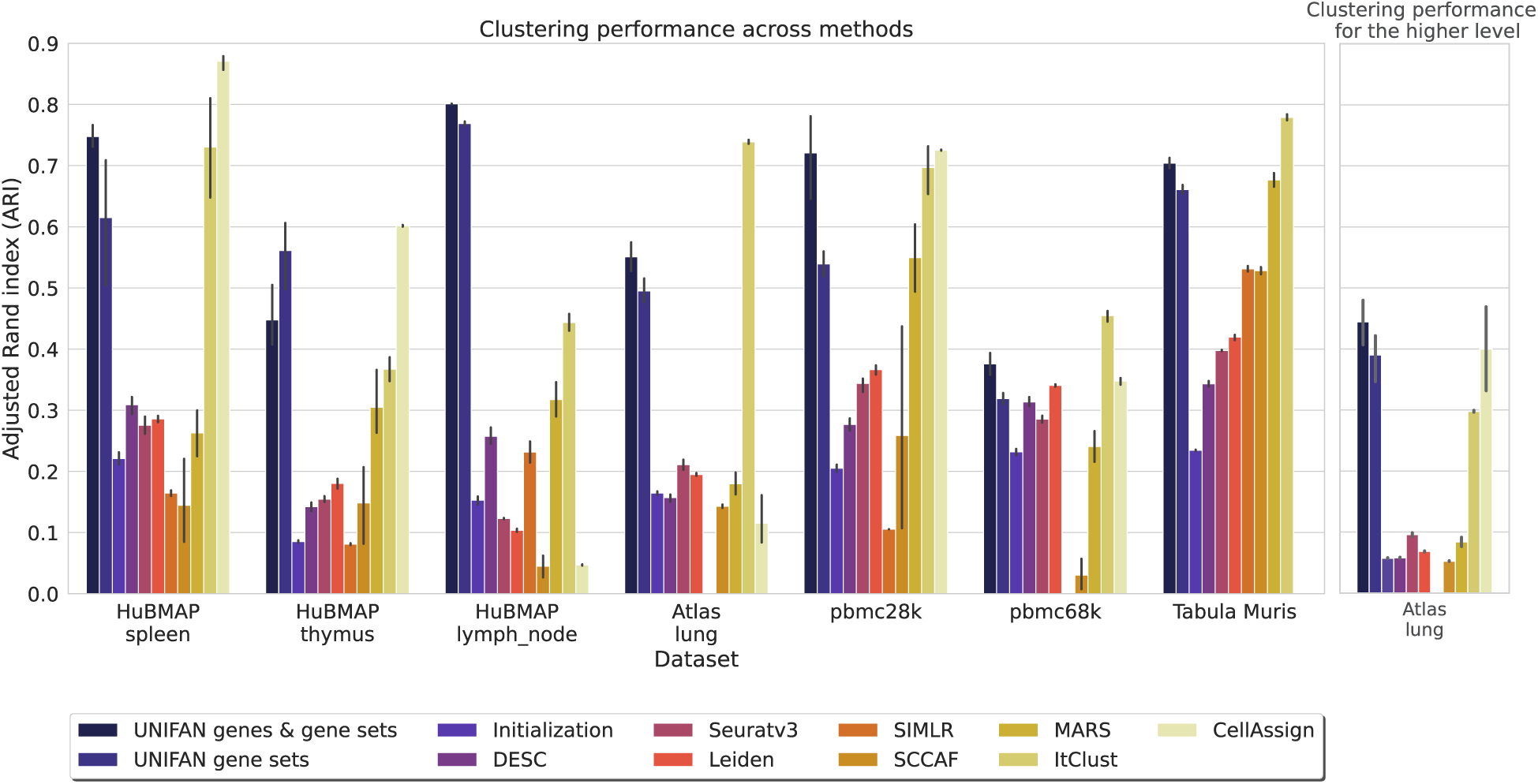
UNIFAN significantly outperforms other methods. “UNIFAN genes & gene sets” is the default UNIFAN version using both gene set activity scores and a subset of genes as features for the annotator. “UNIFAN gene sets” uses only the gene set activity scores. “Initialization” is the initialization clustering results. The others are the prior methods we used for comparison. For the Tabula Muris data, we take the average over all tissues. See Figure S11 and S12 for tissue specific results. The “Atlas lung” data provides two levels of cell type annotations and so we show results for both (less detailed annotation comparison shown on the right). SIMLR was unable to cluster the “pbmc68k” and “Atlas lung” data since it ran out of memory. CellAssign does not have an average over all tissues for “Tabula Muris” because some of the tissues in that dataset do not have matched cell type marker genes. See Supplementary Methods for details.

As for the (semi-)supervised methods, UNIFAN outperforms MARS in all datasets and improves on ItClust for most of them as well (e.g., average ARI/NMI of UNIFAN and the best performing (semi-)supervised prior method on “HuBMAP thymus”: UNIFAN-0.45/0.55, ItClust-0.37/0.46; on “HuBMAP lymph_node”: UNIFAN-0.80/0.71, ItClust-0.44/0.48; on “pbmc28k”: UNIFAN-0.72/0.74, ItClust-0.70/0.70). UNIFAN is worse than ItClust for “pbmc68k” (average ARI/NMI: UNIFAN-0.38/0.54, ItClust-0.45/0.57) and “Atlas lung” (average ARI/NMI: UNIFAN-0.55/0.69, ItClust-0.74/0.78). However, even for “Atlas lung”, UNIFAN improves on ItClust when using another, likely more robust, level of cell annotations (6 general cell types vs. 38 used in the initial comparison) as shown on the right of Figures 3 and S9. For two other datasets, the performance is comparable (average ARI/NMI on “HuBMAP spleen”: UNIFAN-0.75/0.71, ItClust-0.73/0.75; on “Tabula Muris”: UNIFAN-0.70/0.75, ItClust-0.78/0.74).

For the CellAssign comparison, UNIFAN outperforms CellAssign on most datasets (average ARI/NMI on “HuBMAP lymph_node”: UNIFAN-0.80/0.71, CellAssign-0.05/0.15; on “Atlas lung”: UNIFAN-0.55/0.69, CellAssign-0.12/0.24; on “pbmc68k”: UNIFAN-0.38/0.54, CellAssign-0.35/0.53). CellAssign was unable to annotate some tissues in the Tabula Muris data which do not have matched cell type markers in the database (e.g., adipose tissues). For those having matched markers, UNIFAN performs better than CellAssign in the majority of them, as shown in Figures S11 - S14. UNIFAN is worse than CellAssign for “HuBMAP spleen” (average ARI/NMI: UNIFAN-0.75/0.71, CellAssign-0.87/0.72) and “HuBMAP thymus” (average ARI/NMI: UNIFAN-0.45/0.55, CellAssign-0.60/0.56), for both of which the markers of many cell types are well documented in the marker database. For “pbmc28k”, it is hard to tell which method performs better given the disagreements between ARI and NMI (average ARI/NMI: UNIFAN-0.72/0.74, CellAssign-0.73/0.66).

Overall, these results indicate that UNIFAN provides more accurate cell type identification results especially in cases where true cell type labels or cell type markers are not available or very few are known.

To further evaluate the different parts of UNIFAN in order to determine which input or processing is contributing the most to its success, we compared different versions of UNIFAN. These included “UNIFAN no annotator” which is composed of only the clustering part without the annotator, “UNIFAN random” which uses randomly generated features for the annotator and several other variations differing in the biological features used by the annotator including “UNIFAN gene sets” which uses only gene set activity scores and “UNIFAN genes & gene sets” (the default version) which uses both gene set activity scores and the selected genes.

As shown in Figure 3 and S10, the two “UNIFAN” variations using gene sets constantly outperformed the other versions which either did not use an annotator or used randomly generated values as features for the annotator. These results indicate that the use of the annotator to focus on the relevant co-expressed sets of genes is crucial to the performance of UNIFAN.

### UNIFAN Identifies Novel Cell Subtypes

We tested the ability of UNIFAN to identify novel cell subtypes. For this, we looked at clusters identified by UNIFAN that are combined in the original annotations. Such clusters represent unique cell subtypes according to UNIFAN while in the original analysis all cells in these clusters are assigned to the same cell type (see Figure 4 for an example). We found a number of such cases and have looked at their biological relevance.

**Figure 4:**
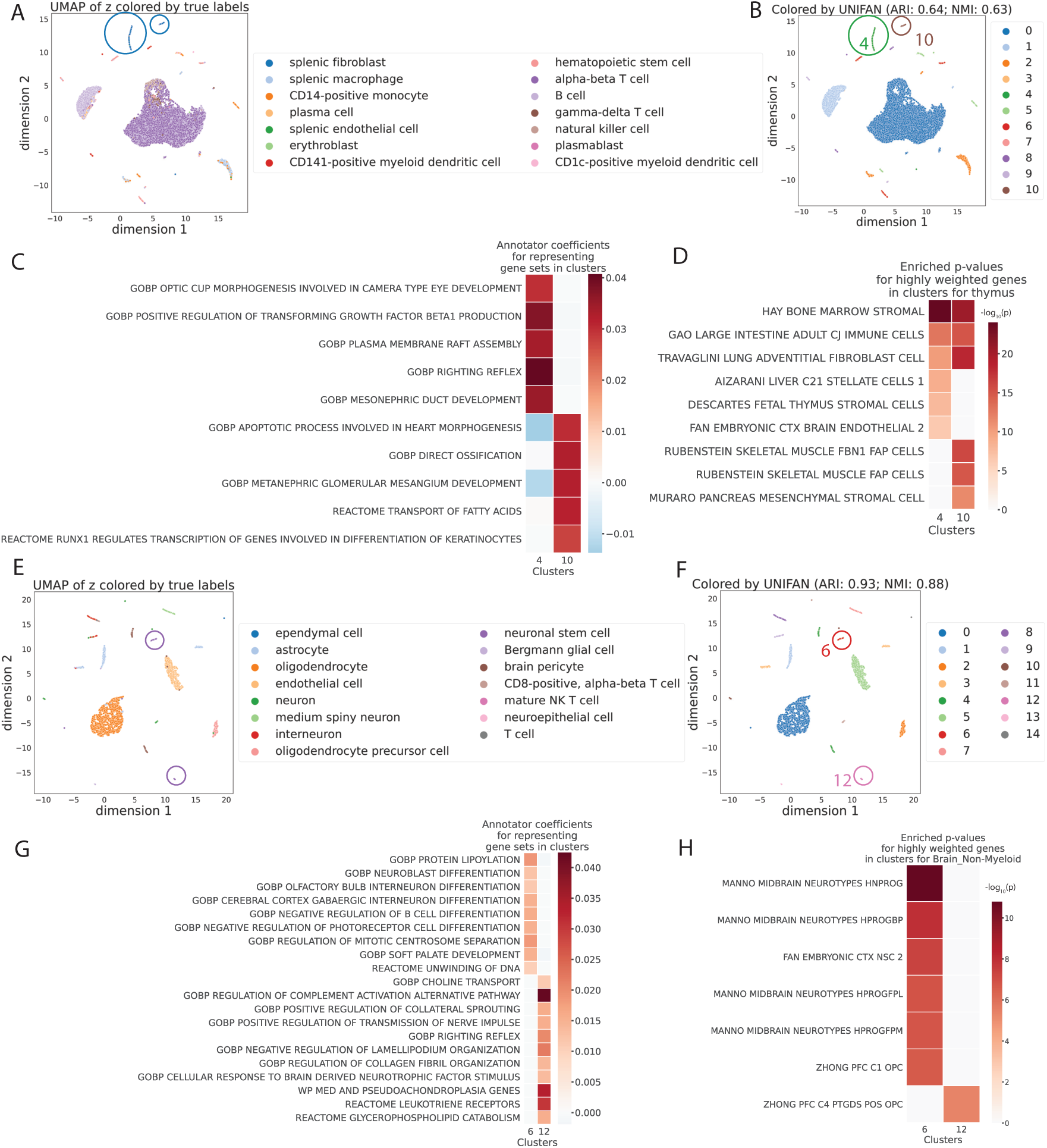
UNIFAN identifies novel cell subtypes for the HuBMAP thymus data and the Brain_Non-Myeloid data in Tabula Muris. **A, B, C and D:** results for the HuBMAP thymus data. **E, F, G, H:** results for the Brain_Non-Myeloid data in Tabula Muris. **A, B, E, F**: UMAP visualization of the low-dimensional representation **z**_**e**_ of cells from UNIFAN. Cluster 4 and 10 are circled for the HuBMAP thymus data. Cluster 6 and 12 are circled for the Brain_Non-Myeloid data in Tabula Muris. **A, E**: Colored by original cell type labels; **B, F**: colored by the clusters identified by UNIFAN. **C, G**: Coefficients learned by the annotator for highly ranked gene sets for the two clusters. **D, H**: Enrichment p-values of cell type marker sets in the highly weighted genes learned by the annotator.

The first example is from UNIFAN’s results of the HuBMAP thymus dataset. Figure 4A and 4B shows a UMAP (Becht et al. 2019) visualization of **z**_**e**_ output from UNIFAN for each cell, colored by the true cell types and clusters found by UNIFAN, respectively. UNIFAN clusters 4 and 10 (circled in Figure 4A and 4B) were both initially labeled as “splenic fibroblast”. Gene sets and markers selected by UNIFAN for both clusters (Figure 4C and 4D) agree with the initial assignment and include “TRAVAGLINI LUNG ADVENTITIAL FIBROBLAST CELL”. They differ, however, in other selected gene sets and markers indicating that they may represent different subtypes. Cluster 4 cells are enriched for thymus stromal, stellate and endothelial cell markers while cluster 10 cells are enriched for markers related to mesenchymal stromal cells (MSCs) including “RUBENSTEIN SKELETAL MUSCLE FBN1 FAP CELLS”, and “MURARO PANCREAS MESENCHYMAL STROMAL CELL” (FAP cells, short form for fibro/adipogenic progenitors, are also a type of MSCs (Wosczyna et al. 2019).).

Examining UNIFAN’s results for the Brain_Non-Myeloid dataset from Tabula Muris (TM Consortium et al. 2018) highlights another novel subtype. In this dataset, we looked at cells labeled as “neuronal stem cell” (shown by the circled clusters in Figure 4E and 4F). UNIFAN assigns these cells to two clusters, 6 and 12. Cluster 6 seems to indeed contain mostly neuronal stem cells since gene sets selected for it are related to the differentiation processes of multiple different cell types (for example, “GOBP NEUROBLAST DIFFERENTIATION” and “GOBP CEREBRAL CORTEX GABAERGIC INTERNEURON DIFFERENTIATION”. See Figure 4G.), indicating its multipotency. The marker set “FAN EMBRYONIC CTX NSC 2” (NSC is the abbreviation for neuronal stem cells) is also enriched in the selected genes (Figure 4H). Cluster 12, however, is likely related to oligodendrocyte precursor cells (OPCs) as the selected genes for this cluster are enriched for “ZHONG PFC C4 PTGDS POS OPC” (Figure 4H). Gene sets related to oligodendrocyte functions including “GOBP POSITIVE REGULATION OF TRANSMISSION OF NERVE PULSE” (Fields 2008) are also selected by UNIFAN for this cluster (Figure 4G).

We also conducted simulation experiments to investigate if UNIFAN is robust when the data contains novel cell types using novel pathways that have not been included in the pathway database used by UNIFAN. Results, presented in Figure S18 show that UNIFAN accurately identities the correct cell types for such simulated data and is able to identify several of the pathways used by the simulated cell types. See Supplementary Results for details.

### Models are Transferable across Tissues and Species

Since different tissues from the same species or the same tissue across species may share cell types, we next explored if an autoencoder for gene set activity scores which is pre-trained on one tissue / species can also be useful for another tissue/species. The importance of such pre-training is that training of the autoencoder for gene set activity scores of UNIFAN is time consuming and so if this can be done offline (i.e. using prior data), then the application of the method to a new dataset can be much faster.

For this, we pre-train a gene set activity scores model using all available human datasets and apply the learned model to infer the gene set activity scores for Tabula Muris mouse datasets. We then run the clustering and annotation using these inferred scores and compare the results with those inferred from a model that was directly trained on the Tabula Muris data as discussed above. Given we focus on the usefulness of gene set activity scores, we use only these scores as features for the annotator (“UNIFAN gene sets”) for this comparison.

Figure 5 presents the results. As expected, we see an overall decline in the average performance over tissues when comparing the results of pre-trained and de novo models. However, for those mouse tissues that are also profiled in the human datasets we used, we observe similar performance when using the pre-trained human model. This is most apparent for spleen, lung and a few adipose tissues including SCAT (subcutaneous adipose tissue) and GAT (gonadal adipose tissue), as shown in Figure 5 and Figure S15-S16. These adipose tissues contain many immune cell types which are also present in many of the human tissues we used for pre-training (spleen, thymus, lymph nodes and PBMC). We further tested pre-trained models for individual tissues (i.e., training using spleen in human and testing only on spleen in mouse). As shown in Figure 5, for such analysis the performance is even better for the most part when compared to using the generally trained model. The only exception is thymus, where the mouse and human annotations differ significantly in the datasets we used. The major cell type in the Tabula Muris thymus data (thymocyte) does not appear in the HuBMAP human thymus data.

**Figure 5:**
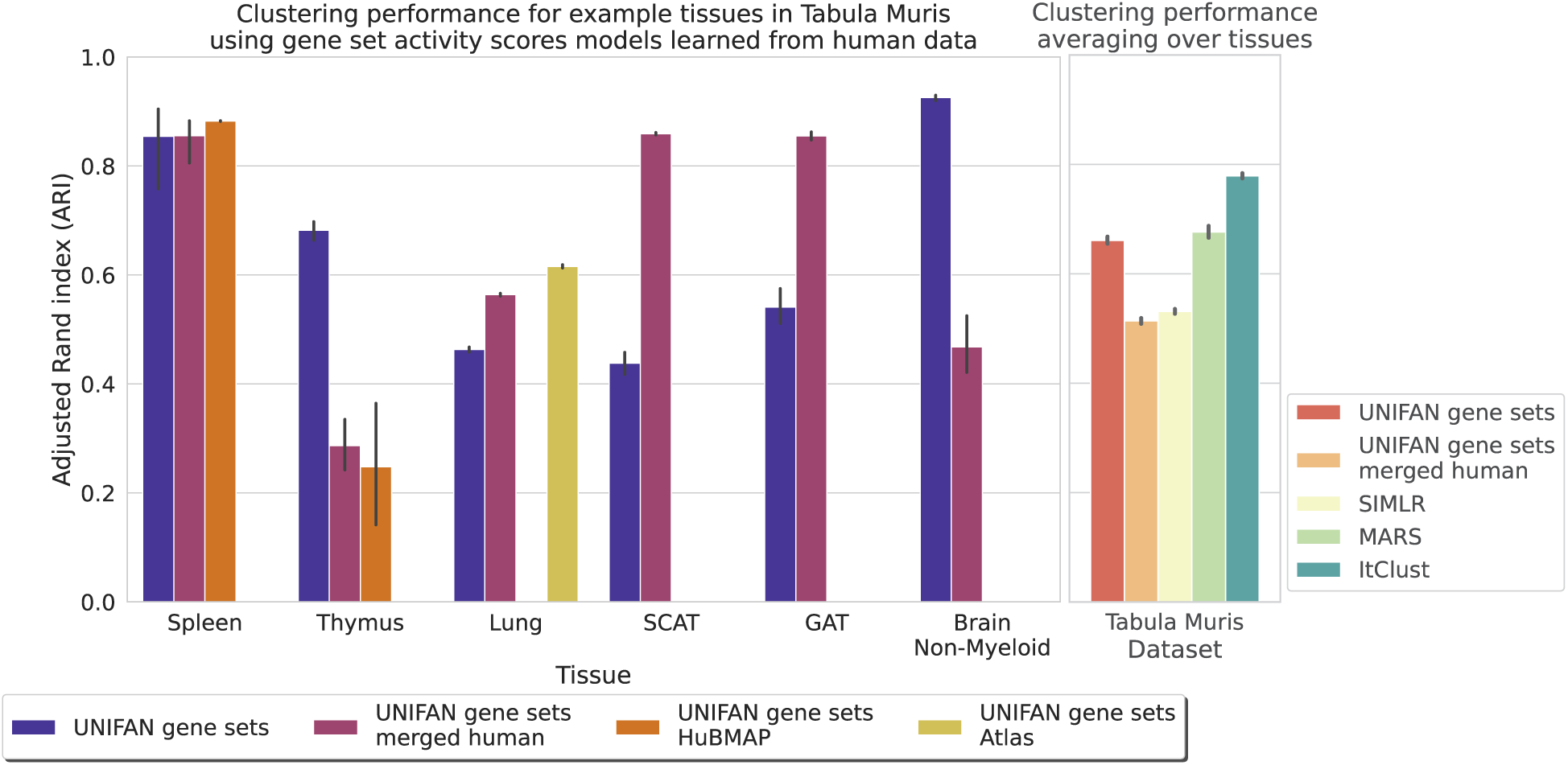
Performance on the Tabula Muris data when using gene set activity scores models pre-trained on human tissues. The plot on the left shows the ARI for some example tissues and the one on the right shows the average ARI across tissues. All versions of UNIFAN methods use models pre-trained on human tissues except for “UNIFAN gene sets”, which used models trained on the same datasets as we discussed before. “UNIFAN gene sets merged human” uses the model pre-trained on all available human tissues. “UNIFAN gene sets HuBMAP” uses the model pre-trained on the corresponding HuBMAP tissue (HuBMAP spleen or thymus). “UNIFAN gene sets Atlas” uses the model pre-trained on the “Atlas lung” dataset. We included only the best performing prior methods on the Tabula Muris data (SIMLR, MARS, ItClust) for comparison. We see that the model pre-trained on human data helpful for mouse gene set activity scores inference and for clustering, specifically for tissues having similar cell types between human and mouse such as spleen and lung. For thymus and brain non-myeloid whose cell types are not well represented in the pre-training set, the performance drops.

## Discussion

Cell type assignment is one of the most important steps in scRNA-Seq analysis. In most cases, such assignment is performed by first clustering cells and then assigning each cluster with a cell type based on differentially expressed genes or the expression of known cell type markers.

Here we presented UNIFAN which improves both clustering and cluster annotations by using a large collection of gene sets (Subramanian et al. 2005). UNIFAN infers gene set activity scores and uses them to regularize the clustering of cells. Such design improves the ability to identify biologically meaningful co-expressed genes and to use these to group cells. In addition to leading to improved clustering, UNIFAN also assigns a subset of the gene sets to clusters which can help characterize their cell types. Finally, by relying on known functional gene sets UNIFAN can further refine cell subtype assignment allowing it to obtain a better resolution for cell type assignments. As we have shown for both human and mouse datasets, such analysis can lead to the identification of novel cell subtypes which may be important as more scRNA-Seq data accumulates.

We compared UNIFAN to several popular methods for clustering scRNA-Seq data using datasets spanning a large number of organs from both human and mouse. As we show, UNIFAN consistently outperforms other methods across these datasets. We also compared UNIFAN with two (semi-) supervised methods for cell type identification (MARS (Brbić et al. 2020) and ItClust (Hu et al. 2020)) and a method based on known cell type markers (CellAssign (AW Zhang et al. 2019)). The semi-supervised methods require at least some labeled data while CellAssign uses as input known cell type markers. Still, as we show in Figures 3 and S9, UNIFAN outperforms these methods for most datasets. We also observed that, for CellAssign and ItClust, when very high quality marker sets and labels were available for specific tissues, the performance of these two methods was better than UNIFAN. Thus, we conclude that while UNIFAN is better for general use, in cases where high confidence marker lists or labels are available for a certain tissue in a certain species methods using labels or markers such as ItClust or CellAssign may be better. We also analyzed the gene sets selected by UNIFAN for various clusters and demonstrated that they match well with the known cell types.

Analysis of the various parts of UNIFAN identified the annotator and the gene sets and genes it uses as the main sources for the improvement. The fact that adding variable genes as input improves performance is likely the result of the fact that current gene sets, while very useful, are incomplete. It is likely that we are still missing from current collections sets of genes characterizing some less known biological processes. In such cases, the selected genes capture groupings that are missed by the known gene sets.

UNIFAN can be slow on large datasets (Table S4 in Supplementary Results). The main time consuming part is training the gene set activity scores model for the data being clustered. To speed up the analysis, we have applied UNIFAN to a new dataset using a gene set activity scores model pre-trained on another dataset. This greatly reduced run time (Supplementary Results) but did lead to drop in performance for tissues whose cell types were not well-represented in the pre-training dataset. As we obtain more data from tissues and conditions, we expect that we can further improve the ability to use pre-training to improve runtime.

UNIFAN is written in Python using PyTorch (Paszke et al. 2017) and is freely avialable from https://github.com/doraadong/UNIFAN.

## Methods

### Datasets and Data Preprocessing

We used both human and mouse datasets from several tissues to test our method. The human samples include three scRNA-Seq datasets from The Human BioMolecular Atlas Program (HuBMAP) consortium (H Consortium et al. 2019). These include “HuBMAP spleen”, “HuBMAP thymus” and “HuBMAP lymph_node”. We use Scanpy (Wolf et al. 2018) for the data preprocessing leading to 34,515 cells and 26,092 genes for “HuBMAP spleen”, 22,367 cells and 24,396 genes for “HuBMAP thymus”, and 24,311 cells and 20,946 genes for “HuBMAP lymph_node”. The “Atlas lung” uses the healthy control samples from Adams et al. (2020). After filtering, this dataset is composed of 96,282 cells and 17,315 genes. The “pbmc28k” data is from Van Der Wijst et al. (2018) and has 25,185 cells and 19,404 genes. The “pbmc68k” data is from Zheng et al. (2017), having 68,551 cells and 17,788 genes. Mouse datasets are from the Tabula Muris paper (TM Consortium et al. 2018). Following Brbić et al. (2020), we end up with 21 datasets each for a single tissue. They all have 22,904 genes and the number of cells ranges from 366 (Aorta) to 4,433 (Heart). See Supplementary Methods for the preprocessing details.

In addition to expression data, UNIFAN uses gene sets to guide clustering. For this we use 7481 gene sets derived from the GO Biological Process ontology (termed c5.go.bp in MSigDB (Subramanian et al. 2005)), 2922 gene sets from pathway databases (c2.cp in MSigDB (Subramanian et al. 2005)) and 335 sets of targets of transcription factors from Ernst et al. (2007). Names for biological process sets start with “GOBP”. Pathway sets use a prefix representing the pathway database they are extracted from (e.g., “KEGG”, “WP”, “REACTOME”). We purposely did not use cell type marker gene sets (c8.all in MSigDB) since we wanted to keep the method unsupervised and marker lists are often based on DE analysis of labeled cell type data.

### Clustering and Annotating Single Cells Using Gene Sets

To enable the use of prior knowledge on gene function and regulation for clustering single cells, we developed a deep learning model, UNIFAN (**Un**supervised S**i**ngle-cell **F**unctional **An**notation). For each single cell, UNIFAN first infers gene set activity scores associated with this cell using the input gene sets. Next, UNIFAN clusters cells by using the learned gene set activity scores and a reduced dimension representation of the expression of genes in the cell. The gene set activity scores are used by an “annotator” to guide the clustering such that cells sharing similar biological processes are more likely to be grouped together. Such design allows the method to focus on the key processes when clustering cells and so can overcome issues related to noise and dropout while simultaneously selecting marker gene sets which can be used to annotate clusters.

### Learning Gene Set Activity Scores for Cells

For each cell, we first infer its gene set activity scores 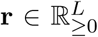 (*L*: number of gene sets), which represent the activity of known biological processes or pathways in the cell. For this, we design a special autoencoder whose decoder, instead of being fully-connected, is composed of a binary matrix *D* ∈ ℝ^*G*×*L*^, where *G* is the total number of genes profiled. Each column in *D* corresponds to a known gene set for a biological process or pathway where the values are indicators for whether a gene belongs to this set or not. For a cell with expression **y**, the encoder, which composes of fully-connected layers, outputs a low-dimensional representation **r. r** is then multiplied by the binary matrix *D* which leads to a reconstructed expression vector **ŷ**, as shown in Figure 6. Values in **r** serve as weights / coefficients for known gene sets for this cell. Parameters for the fully-connected encoder are optimized such that the combination of the gene sets, weighted by **r**, can be used to reconstruct the observed expression **y** for all genes in the cell. Thus, **r** can be seen as the activity levels of pathways and processes in the cell.

**Figure 6:**
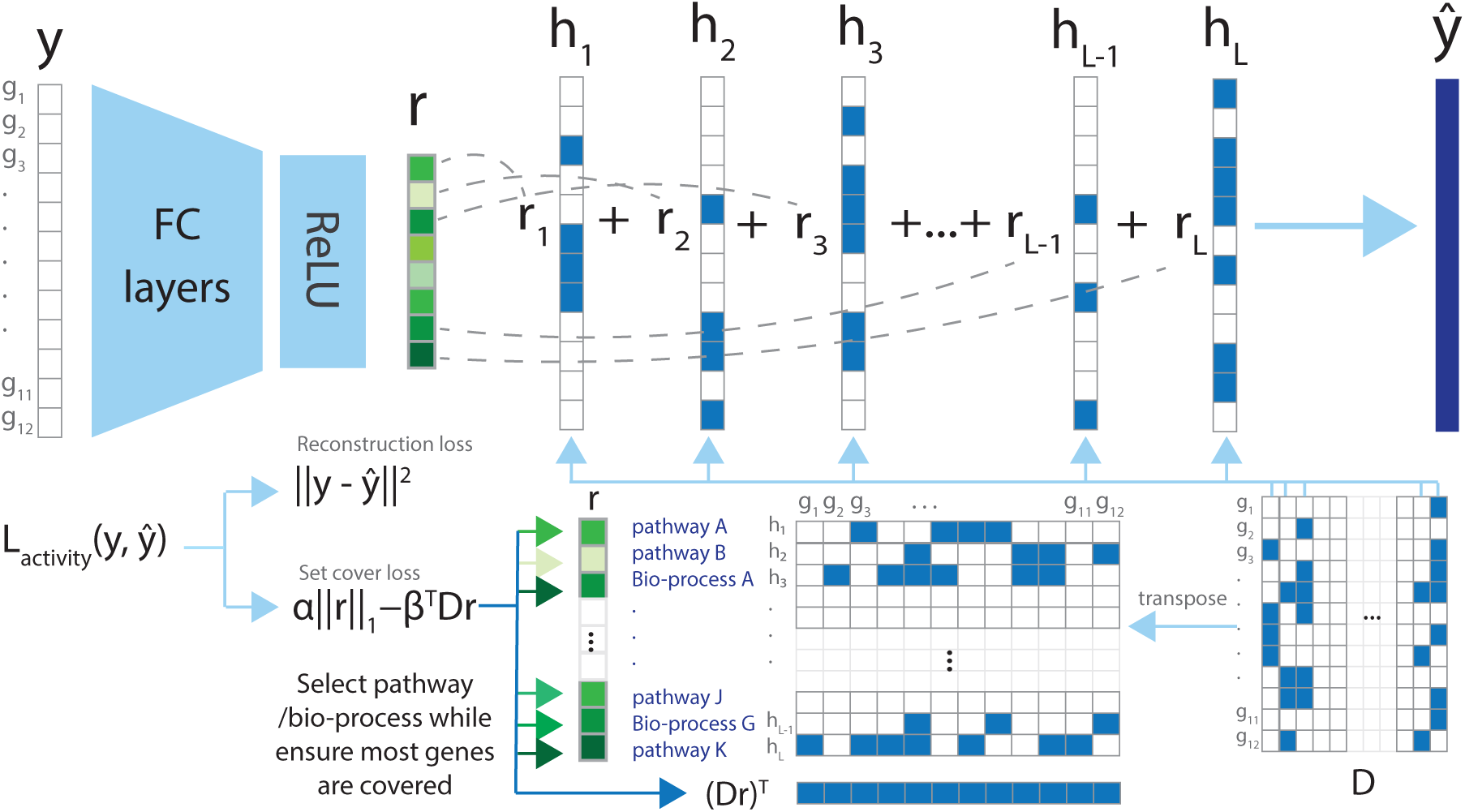
Assigning single-cell gene set activity scores using an autoencoder. The autoencoder is designed such that the decoder is composed of binary vectors with values indicating if a profiled gene belongs to a known gene set or not. The output of the encoder, **r**, serve as coefficients for the gene set vectors, showing how related a cell is to a known pathway/biological processes. **r** thus can be seen as the gene set activity scores for this cell. The set cover loss is designed to select uncorrelated pathways/processes to better annotate cells. FC layers: fully-connected layers. Bio-process: biological process

To construct the gene set matrix *D* which serves as an input to UNIFAN, we collected gene sets representing biological processes (including canonical pathways and targets of specific regulators) from MSigDB (Subramanian et al. 2005) and Ernst et al. (2007), which resulted in a total of roughly 10K sets. We expect that only a small subset of these biological processes are active for each single cell and so we employ regularization to select active gene sets for each cell. First, we constrain **r** to be non-negative by using ReLU for the output layer, which results in most values in **r** being assigned to 0. Next, we use a regularizer inspired by the classical set cover algorithm, which aims to find the least number of sets that cover all elements (in our case, profiled genes of the cell). By employing this regularizer, we aim to find a small subset of non-overlapping gene sets that can cover as many of genes as possible (Lu et al. 2008). For this, our regularizer optimizes the following function *α* ‖ **r**‖_1_ − *β* ^*T*^ *D***r**, where *α* and *β* are hyperparameters (see section Training UNIFAN and Hyperparameter Selection on selecting values for hyperparameters in our model). Using mean-squared error for the reconstruction loss, our overall loss function for a single cell is

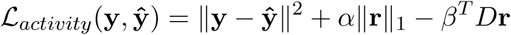

### Clustering Cells Using Gene Set Activity Scores

To cluster cells using the inferred gene set activity scores, we use an autoencoder-based method. It is composed of two parts, an expression based cluster assignment part (“grey” parts in Figure 7) and the “annotator” part (“green” parts in Figure 7) which uses the gene set activity scores discussed above as input.

**Figure 7:**
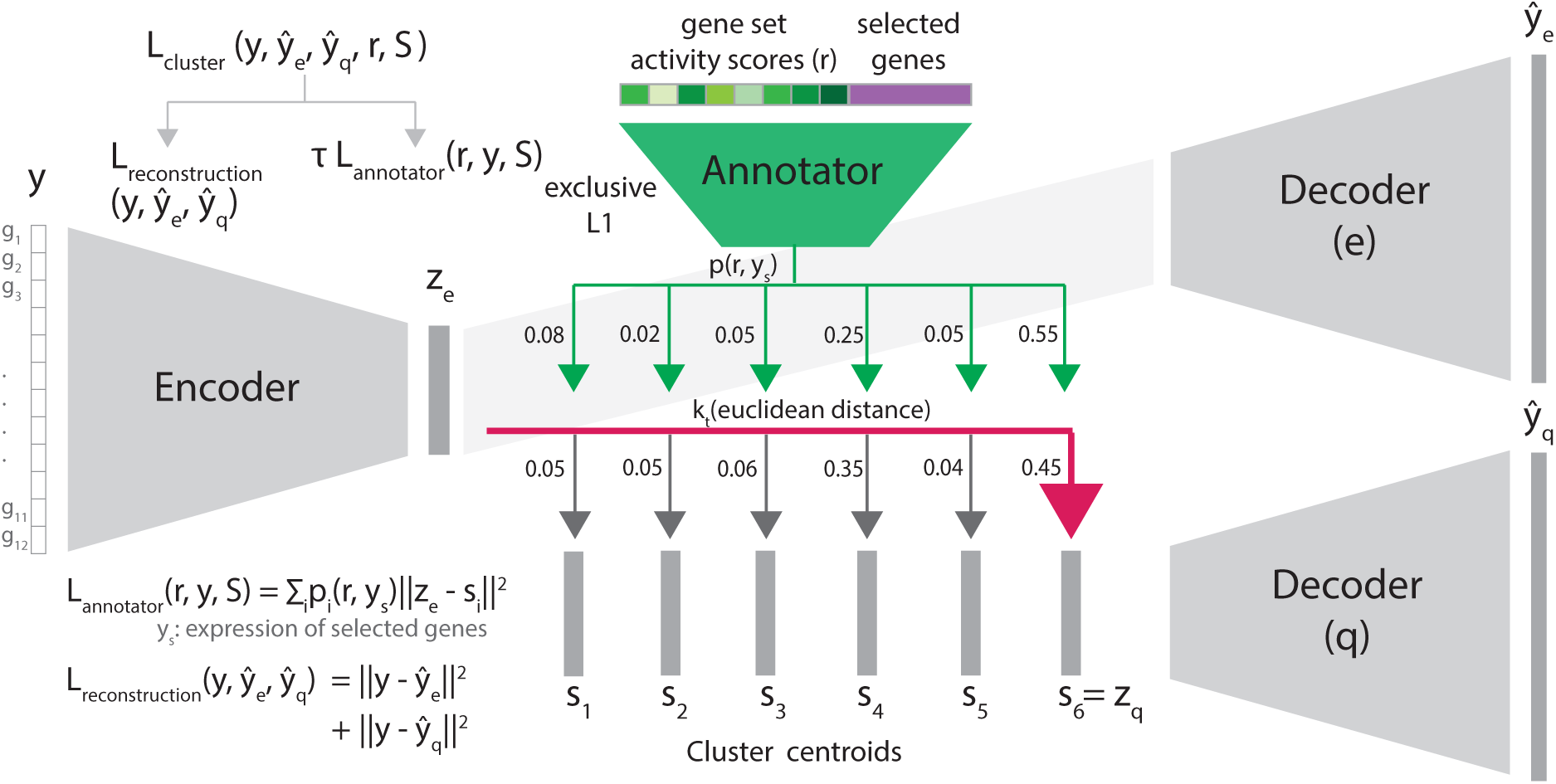
Jointly clustering and annotating cells. The autoencoder contains two parts: the cluster assignment part (grey in Figure 7) uses a low-dimensional representation **z**_**e**_ to assign a cell to clusters; the “annotator” (green in Figure 7) uses the learned gene set activity scores and selected genes’ expression to refine clustering and annotate clusters. Gene sets and genes selected as predictive by the annotator, in turn, provide useful annotations for each cell cluster. We set the number of clusters M as 6 in this figure for illustration purposes.

The cluster assignment part only uses the expression profile for each cell. It consists of an encoder and two decoders (Decoder(e) and (q) in Figure 7), modified based on Fortuin et al. (2018) and Oord et al. (2017). For a single cell, we first use an encoder on the expression of genes in the cell **y**, resulting in a low-dimensional representation **z**_**e**_, as shown in Figure 7. After initialization, we start with a guess of *M* clusters and cluster centroids. Among *M* cluster centroids *S* = *{***s**_**1**_, **s**_**2**_, …, **s**_**M**_*}*, we identify the centroid closest to **z**_**e**_ by first calculating the euclidean distances between **z**_**e**_ and all centroids and then transforming the distances using a t-distribution kernel 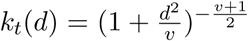, following Li et al. (2020), Xie et al. (2016), and van der Maaten and Hinton (2008). *d* stands for the distance and *v* stands for the degrees of freedom, which is fixed at 10 for all experiments. We then take the closest centroid as the discrete representation **z**_**q**_ of the cell and assign the cell to the corresponding cluster. We obtain the reconstructed expression **ŷ**_**q**_ using decoder (q) which only takes **z**_**q**_ as input and so all cells in the same cluster have the same reconstructed expression. We optimize the reconstruction error ‖**y** *-* **ŷ**_**q**_‖^2^ to find the best **z**_**q**_, cluster centroids *S*, and decoder (q), in a manner similar to finding the best cluster centroids in k-means clustering.

Since we assign cell clusters using k-means (i.e. discrete assignment), the encoder cannot be learned using backpropagation. To enable the iterative refinement of model parameters using gradients, we follow Fortuin et al. (2018) by adding another decoder, decoder (e). Decoder (e) takes **z**_**e**_ as input and outputs another reconstructed expression **ŷ**_**e**_. By optimizing ‖**y** *−* **ŷ**_**e**_‖^2^, we can now update **z**_**e**_ and the encoder. The overall loss function for a single cell in the cluster assignment part is thus ℒ_reconstruction_(**y, ŷ**_**e**_, **ŷ**_**q**_)= ‖**y** − **ŷ**_**e**_‖^2^ + ‖**y** − **ŷ**_**q**_‖^2^. All neural networks mentioned above are composed of fully-connected layers.

So far we only discussed clustering using expression data. We next use the learned gene set activity scores for each cell to refine cluster assignments as well as to annotate cell clusters. For this, we add an “annotator”, a logistic classifier, to the network model. For each cell, the annotator uses the gene set activity scores **r** for that cell as input and outputs **p**(**r**), the probability of the cell being assigned to each cluster. We use the annotator’s output to refine the cluster assignment by adding 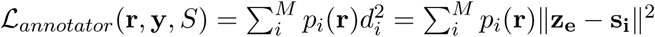 to the existing loss function. Since it uses both, the low-dimensional representation of the cell **z**_**e**_ and the cluster centroids S, such loss encourages cells being assigned to clusters based on the probability specified by **p**(**r**). In other words, by employing ℒ_*annotator*_ and the annotator, we are using prior knowledge about gene membership in key biological processes to guide the dimension reduction and cluster assignment. Gene sets selected as predictive by the annotator, in turn, provide useful annotations for each cell cluster.

To allow the selection of marker genes for each cluster, we also tested the use of the UNIFAN with a subset of the most variable genes selected using Seuratv3 (Stuart et al. 2019). Using such set the annotator loss becomes: 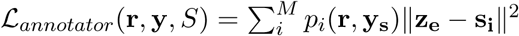, where **y**_**s**_ are the expression of the selected genes. The overall loss function for the cluster assignment part is thus

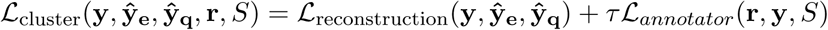

where *τ* is a weighting hyperparameter.

The annotator is trained to optimize its own loss function. We use cross-entropy loss to train the annotator: 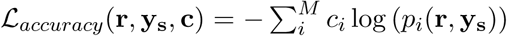, where *c*_*i*_ =𝟙 (cell clustered to *i*). To select marker gene sets and genes specific to each cluster, we use the exclusive LASSO regularizer (Zhou et al. 2010) for the annotator. The regularizer takes the form of 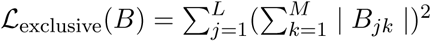, where *B* are the parameters of the logistic classifier. Thus, the overall loss function for the annotator is

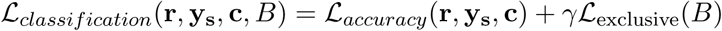

where *γ* is a weighting hyperparameter.

### Training UNIFAN and Hyperparameter Selection

The loss functions described in the section above are defined for a single cell. During training, we take the mean of loss over all cells. We first train the autoencoder for the gene set activity scores and obtain the gene set activity scores for all cells. We then pre-train the encoder and the decoder (e) of the autoencoder for clustering on the expression data to obtain an initial low-dimensional representations of the cells. We then run Leiden clustering (Traag et al. 2019) on these representations to obtain a guess of the number of clusters *M*, the initial cluster assignment and the cluster centroids *S*. Both the number of clusters *M* and cluster centroids *S* are refined later as part of the training. Specifically, clusters with no cell assigned to them are removed. The annotator is then pre-trained using the inferred gene set activity scores and the selected genes, if available. We use the cluster assignment initialized as described above as the true labels to pre-train the annotator.

Finally, we train the annotator together with the cluster assignment part (the encoder and decoder (e) & decoder(q)). In each epoch, the annotator is trained by using the clustering results as the true label for each cell. The output from the annotator **p**(**r**) is in turn used to evaluate the annotator loss ℒ_*annotator*_ for the cluster assignment part. As described previously, the annotator is optimized using its own loss function, separately from the cluster assignment part. See Supplementary Methods for details in training.

We use 32 dimensions for the low-dimensional representation **z**_**e**_ of a cell. To select the values for hyperparameters, including the neural network configuration and the weighting hyperparameters for the loss functions, we conducted a grid search using the Tabula Muris dataset and selected those hyperparameters values leading to the best performance over tissues. Unless specifically mentioned, the same set of values were applied to all datasets in all experiments. See Supplementary Methods for details on how we select the values. As discussed in Supplementary Results, our method is robust to different choices of hyperparameter values. We also show in Supplementary Results that our method is robust to the resolution value used in Leiden clustering initialization and to the number of clustering epochs.

### Performance Evaluation and Comparison to Other Methods

To evaluate the performance of UNIFAN and to compare it to prior methods including Leiden clustering (Traag et al. 2019), Seuratv3 (Stuart et al. 2019), SIMLR (Wang et al. 2017), DESC (Li et al. 2020), SCCAF (Miao et al. 2020), MARS (Brbić et al. 2020), ItClust (Hu et al. 2020) and CellAssign (AW Zhang et al. 2019), we run each method on each dataset ten times with different initializations. For the Tabula Muris data, we run methods on each tissue separately. We use adjusted Rand index (ARI) and Normalized Mutual Information (NMI) implemented in scikit-learn (Pedregosa et al. 2011) to compare clusters with ground truth annotations. Since calculating ARI for large datasets is time consuming, we use stratified random sampling when computing ARI for large datasets (>5e4 cells).

For the two (semi-) supervised methods MARS and ItClust, for each dataset, we use random sampling stratified according to the true cell types to select 5% of the cells for training where the methods take also the true labels as input, following Wei and S Zhang (2021). We leave the rest 95% of the cells for testing and use these to calculate ARI and NMI. We re-sample the cells for training each time we run the two methods. For CellAssign which takes known cell type markers as input, we use the marker gene sets from PanglaoDB (Franzén et al. 2019), as described in AW Zhang et al. (2019). Using all available cell type markers as input for a dataset increases run time dramatically and so we only use cell type markers for the tissue that best matches the tissue of the dataset (for example, we use marker genes for “Immune system” for the dataset “pbmc28k”).

See Supplementary Methods for details on how we used prior methods, including hyper-parameter settings for these methods and for information on the evaluation strategy including how we compute enrichment p-values of the cell type marker sets in the highly weighted genes learned by the annotator. See also Supplementary Methods for details on how we conducted the simulation experiments.

## Supporting information

Supplementary Methods and Results

## Software availability

UNIFAN is available under a MIT license at Github (https://github.com/doraadong/UNIFAN) and as Supplemental Code.

## Acknowledgments

Work partially supported by NIH grants OT2OD026682, 1U54AG075931 and 1U24CA268108 to Z.B.J.

## Author Contributions

D.L., J.D., Z.B.-J. developed the method; D.L. implemented the software and performed the analysis; All authors analyzed the results and wrote the manuscript.

## References

Abdelaal T, Michielsen L, Cats D, Hoogduin D, Mei H, Reinders MJ, and Mahfouz A. 2019. A comparison of automatic cell identification methods for single-cell RNA sequencing data. Genome Biol. 20: 1–19.

Adams TS et al. 2020. Single-cell RNA-seq reveals ectopic and aberrant lung-resident cell populations in idiopathic pulmonary fibrosis. Sci. Adv. 6: eaba1983.

Alavi A, Ruffalo M, Parvangada A, Huang Z, and Bar-Joseph Z. 2018. A web server for comparative analysis of single-cell RNA-seq data. Nat. Commun. 9: 1–11.

Becht E, McInnes L, Healy J, Dutertre CA, Kwok IW, Ng LG, Ginhoux F, and Newell EW. 2019. Dimensionality reduction for visualizing single-cell data using UMAP. Nat. Biotechnol. 37: 38–44.

Blondel VD, Guillaume JL, Lambiotte R, and Lefebvre E. 2008. Fast unfolding of communities in large networks. J. Stat. Mech: Theory Exp. 2008: P10008.

Brbic M, Zitnik M, Wang S, Pisco AO, Altman RB, Darmanis S, and Leskovec J. 2020. MARS: discovering novel cell types across heterogeneous single-cell experiments. Nat. Methods. 17: 1200–1206.

Clarke ZA, Andrews TS, Atif J, Pouyabahar D, Innes BT, MacParland SA, and Bader GD. 2021. Tutorial: guidelines for annotating single-cell transcriptomic maps using automated and manual methods. Nat. Protoc. 16: 2749–2764.

Consortium H et al. 2019. The human body at cellular resolution: the NIH Human Biomolecular Atlas Program. Nature. 574: 187.

Consortium TM et al. 2018. Single-cell transcriptomics of 20 mouse organs creates a Tabula Muris. Nature. 562: 367–372.

Ernst J, Vainas O, Harbison CT, Simon I, and Bar-Joseph Z. 2007. Reconstructing dynamic regulatory maps. Mol. Syst. Biol. 3: 74.

Fields RD. 2008. Oligodendrocytes changing the rules: action potentials in glia and oligodendrocytes controlling action potentials. The Neuroscientist. 14: 540–543.

Fortuin V, Hüser M, Locatello F, Strathmann H, and Rätsch G. 2018. Som-vae: Interpretable discrete representation learning on time series. arXiv preprint 1806.02199.

Franzén O, Gan LM, and Björkegren JL. 2019. PanglaoDB: a web server for exploration of mouse and human single-cell RNA sequencing data. Database. 2019:

Gayoso A, Lopez R, Xing G, Boyeau P, Valiollah Pour Amiri V, Hong J, Wu K, Jayasuriya M, Mehlman E, Langevin M, et al. 2022. A Python library for probabilistic analysis of single-cell omics data. Nat. Biotechnol. 1–4.

Hu J, Li X, Hu G, Lyu Y, Susztak K, and Li M. 2020. Iterative transfer learning with neural network for clustering and cell type classification in single-cell RNA-seq analysis. Nat. Mach. Intell. 2: 607–618.

Kingma DP and Ba J 2014. Adam: A Method for Stochastic Optimization. eprint: 1412.6980.

Kiselev VY, Yiu A, and Hemberg M. 2018. scmap: projection of single-cell RNA-seq data across data sets. Nat. Methods. 15: 359–362.

Li X, Wang K, Lyu Y, Pan H, Zhang J, Stambolian D, Susztak K, Reilly MP, Hu G, and Li M. 2020. Deep learning enables accurate clustering with batch effect removal in single-cell RNA-seq analysis. Nat. Commun. 11: 1–14.

Lu Y, Rosenfeld R, Simon I, Nau GJ, and Bar-Joseph Z. 2008. A probabilistic generative model for GO enrichment analysis. Nucleic Acids Res. 36: e109–e109.

Miao Z, Moreno P, Huang N, Papatheodorou I, Brazma A, and Teichmann SA. 2020. Putative cell type discovery from single-cell gene expression data. Nat. Methods. 17: 621–628.

Oord Avd, Vinyals O, and Kavukcuoglu K. 2017. Neural discrete representation learning. arXiv preprint 1711.00937.

Paszke A, Gross S, Chintala S, Chanan G, Yang E, DeVito Z, Lin Z, Desmaison A, Antiga L, and Lerer A. 2017. Automatic differentiation in PyTorch.

Pedregosa F et al. 2011. Scikit-learn: Machine Learning in Python. J. Mach. Learn. Res. 12: 2825–2830.

Pliner HA, Shendure J, and Trapnell C. 2019. Supervised classification enables rapid annotation of cell atlases. Nat. Methods. 16: 983–986.

Schmidl C, Renner K, Peter K, Eder R, Lassmann T, Balwierz PJ, Itoh M, Nagao-Sato S, Kawaji H, Carninci P, et al. 2014. Transcription and enhancer profiling in human monocyte subsets. Blood. 123: e90–e99.

Stuart T, Butler A, Hoffman P, Hafemeister C, Papalexi E, Mauck III WM, Hao Y, Stoeckius M, Smibert P, and Satija R. 2019. Comprehensive integration of single-cell data. Cell. 177: 1888–1902.

Subramanian A et al. 2005. Gene set enrichment analysis: A knowledge-based approach for interpreting genome-wide expression profiles. Proc. Natl. Acad. Sci. U. S. A. 102: 15545–15550.

Tanay A and Regev A. 2017. Scaling single-cell genomics from phenomenology to mechanism. Nature. 541: 331–338.

Traag VA, Waltman L, and Van Eck NJ. 2019. From Louvain to Leiden: guaranteeing well-connected communities. Sci. Rep. 9: 1–12.

van der Maaten L and Hinton G. 2008. Visualizing High-Dimensional Data Using t-SNE. J. Mach. Learn. Res. 9: 2579–2605.

Van Der Wijst MG, Brugge H, De Vries DH, Deelen P, Swertz MA, and Franke L. 2018. Single-cell RNA sequencing identifies celltype-specific cis-eQTLs and co-expression QTLs. Nat. Genet. 50: 493–497.

Wang B, Zhu J, Pierson E, Ramazzotti D, and Batzoglou S. 2017. Visualization and analysis of single-cell RNA-seq data by kernel-based similarity learning. Nat. Methods. 14: 414–416.

Wei Z and Zhang S. 2021. CALLR: a semi-supervised cell-type annotation method for single-cell RNA sequencing data. Bioinformatics. 37: i51–i58.

Wolf FA, Angerer P, and Theis FJ. 2018. SCANPY: large-scale single-cell gene expression data analysis. Genome Biol. 19: 1–5.

Wosczyna MN, Konishi CT, Carbajal EEP, Wang TT, Walsh RA, Gan Q, Wagner MW, and Rando TA. 2019. Mesenchymal stromal cells are required for regeneration and homeostatic maintenance of skeletal muscle. Cell Rep. 27: 2029–2035.

Xie J, Girshick R, and Farhadi A 2016. Unsupervised deep embedding for clustering analysis. In: International conference on machine learning. PMLR, pp. 478–487.

Xin J, Mark A, Afrasiabi C, Tsueng G, Juchler M, Gopal N, Stupp GS, Putman TE, Ainscough BJ, Griffith OL, et al. 2016. High-performance web services for querying gene and variant annotation. Genome Biol. 17: 1–7.

Zhang AW, O’Flanagan C, Chavez EA, Lim JL, Ceglia N, McPherson A, Wiens M, Walters P, Chan T, Hewitson B, et al. 2019. Probabilistic cell-type assignment of single-cell RNA-seq for tumor microenvironment profiling. Nat. Methods. 16: 1007–1015.

Zheng GX, Terry JM, Belgrader P, Ryvkin P, Bent ZW, Wilson R, Ziraldo SB, Wheeler TD, McDermott GP, Zhu J, et al. 2017. Massively parallel digital transcriptional profiling of single cells. Nat. Commun. 8: 1–12.

Zhou Y, Jin R, and Hoi SCH 2010. Exclusive lasso for multi-task feature selection. In: Proceedings of the thirteenth international conference on artificial intelligence and statistics. JMLR Workshop and Conference Proceedings, pp. 988–995.

