## Supplementary Methods and Results for "Unsupervised cell functional annotation for single-cell RNA-Seq"

### 1 Supplementary Notes

#### 1.1 Supplementary Methods

##### 1.1.1 Details of Data Preprocessing

We use three scRNA-Seq datasets processed and published by The Human BioMolecular Atlas Program (HuBMAP) consortium (H Consortium et al. 2019). These include "HuBMAP spleen", "HuBMAP thymus" and "HuBMAP lymph_node". We use Scanpy (Wolf et al. 2018) for the data pre-processing. For each dataset, we first filter out cells expressing fewer than 200 genes and genes expressed in fewer than 3 cells. We then normalize each cell to 1e4 total read counts. The values are then log transformed and scaled to zero mean and unit variance. We clip the values such that the maximum is ten. We use MyGene.Info (Xin et al. 2016) to convert the ensemble gene IDs to gene symbols. After preprocessing, we have 34,515 cells × 26,092 genes for "HuBMAP spleen", 22,367 cells × 24,396 genes for "HuBMAP thymus" and 24,311 cells × 20,946 genes for "HuBMAP lymph_node".

The "Atlas lung" data is obtained from the Idiopathic Pulmonary Fibrosis (IPF) Cell Atlas (Adams et al. 2020). We use the healthy control samples. Again, we use Scanpy to first filter out cells expressing fewer than 500 genes and genes expressed in fewer than 5 cells. We then normalize each cell to 1e4 total read counts and log transform the values. Given the large number of genes profiled, we take only a subset of genes by selecting the most variable ones using Scanpy's implementation of "Seurat"-style variable genes selection with minimum mean, maximum mean and minimum dispersion set to 0, 1,000 and 0.01, respectively. Finally, we scale the values to zero mean and unit variance and clip those larger than ten. This results in a dataset of 96,282 cells × 17,315 genes. This dataset has two levels of annotations of the cell types and we use both for evaluation.

The "pbmc28k" data is from Van Der Wijst et al. (2018). We download the counts data and annotations from `https://genenetwork.nl/scrna-seq/`. Same as for other datasets, we use Scanpy to filter out cells expressing fewer than 200 genes and genes expressed in fewer than 3 cells. We then normalize each cell to 1e4 total read counts. The values are then log transformed and scaled to zero mean and unit variance. We clip the values such that the maximum is ten. This results in a dataset of 25,185 cells × 19,404 genes. The "pbmc68k" data is from Zheng et al. (2017) and we downloaded the counts data from `https://www.10xgenomics.com/resources/datasets/fresh-68-k-pbm-cs-donor-a-1-standard-1-1-0`. The annotations are downloaded from `https://github.com/10XGenomics/single-cell-3prime-paper/issues/3`. We use Scanpy to filter out cells expressing fewer than 200 genes and genes expressed in fewer than 3 cells. We then normalize each cell to 1e4 total read counts. The values are then log transformed and scaled to zero mean and unit variance. We clip the values such that the maximum is ten. The resulting dataset is of 68,551 cells × 17,788 genes.

All of the datasets mentioned above are from human tissue samples. We also use the Tabula Muris (TM Consortium et al. 2018) dataset from mouse tissue samples. We downloaded the Tabula Muris Senis data from `https://figshare.com/projects/Tabula_Muris_Senis/64982` and used "cell_ontology_class" as cell type labels. We first use Scanpy to filter out cells expressing fewer than 500 genes or 5000 read counts and genes expressed in fewer than 5 cells. Following Brbić et al. (2020), we take the cells from 3 months old samples from the Tabula Muris Senis data to obtain the Tabula Muris data and we also removed the "Marrow" and "Brain_Myeloid" tissues due to lack of expert validation or stable markers to distinguish cell types. For each of the rest of the tissues, we then normalize the cells to 1e4 total read counts. The values are then log transformed, scaled to zero mean and unit variance and truncated at ten. This results in 21 datasets, each for a single tissue. They all have 22,904 genes and having number of cells ranging from 366 (Aorta) to 4,433 (Heart).

For the merged human dataset which we use to train the gene set activity scores model and apply on mouse tissues, we use all the HuBMAP datasets, the "Atlas lung" and the "pbmc28k" dataset.

##### 1.1.2 Details for Model Training and Hyperparameter Selection

Throughout the training process, we use Adam (Kingma and Ba 2014) as the optimizer and fix the learning rate at 5e-4. We use a batch size of 16 for data with sample size <1e4 and 32 otherwise for all steps except for pre-training the annotator, where we use a batch size of 32 for all datasets. We use 32 dimensions for the low-dimensional representation $\mathbf{z_e}$ of a cell. For the other hyperparameters, including the neural network configuration and the weighting hyperparameters in the loss functions, we conducted a grid search using the Tabula Muris dataset and selected those hyperparameters values leading to the best averaging performance over tissues. Unless specifically mentioned, the same set of values were applied to all datasets in all experiments.

###### 1.1.2.1 Training the Gene Set Activity Scores Model

For the encoder of the autoencoder model, we use 5 fully-connected hidden layers of size 128 with ReLU activations. As noted above, we also use a ReLU activation for the output layer. The drop-out rate is set to 0.1 for all layers. As for the decoder, as described before, we use the binary matrix $D$. We fix the number of training epochs at 70 for all experiments except for training the model for the merged human tissues dataset. Given the large sample size (about 2e5) of the merged data, we train the model for 100 epochs.

There are two weighting hyperparameters in our loss function $\mathcal{L}_{activity}(\mathbf{y}, \hat{\mathbf{y}}) = \|\mathbf{y} - \hat{\mathbf{y}}\|^2 + \alpha\|\mathbf{r}\|_1 - \beta^T D\mathbf{r}$: $\alpha$ and $\beta$. These are used to weight the L1 penalty and the set cover term, respectively. We evaluated how these two hyperparameters impact the sparsity of the resulting gene sets and gene coverage. We quantify the sparsity by the number of non-zero values in the gene set activity scores $\mathbf{r}$ and gene coverage by the number of genes a cell uses and the average number of times a gene is used. As we show in Figure S1, both $\alpha$ and $\beta$ impact the sparsity of the gene set activity scores $\mathbf{r}$ while only $\beta$ impacts the gene coverage, as expected.

Note that $\beta$ is a vector with each element corresponding to a gene. One may apply different values for different genes if weights are available. For our case, we use the same value for all genes. We fix $\alpha$ at 0.01 and $\beta$ at 1e-05.

###### 1.1.2.2 Training the Autoencoder for Cluster Assignment and Annotation

For the encoder, we use 3 fully-connected hidden layers of size 128 with ReLU activation and linear activation for the output layer. We use the same configuration for decoder (e) and (q), both with 2 fully-connected hidden layers of size 128 with ReLU activation. We also use linear activation for the output layers. The drop-out rate is set to 0.1 for all layers. For the annotator, as noted before, we use a logistic regression.

We first pre-train the encoder and the decoder (e) on the expression data for 50 epochs. We use this pre-trained autoencoder to generate initial low-dimensional representations of the cells. We then run Leiden clustering (Traag et al. 2019) implemented in Scanpy (Wolf et al. 2018) on these initial low-dimensional representations to obtain a guess of the number of clusters $M$, the initial cluster assignment and the cluster centroids $S$. We use the default resolution 1. Both the number of clusters $M$ and cluster centroids $S$ are refined as part of the training. Specifically, clusters with no cell assigned to them are removed.

We then pre-train the annotator using the inferred gene set activity scores and the selected genes, if available. The genes are selected using the variable genes selection method of Seuratv3 (Stuart et al. 2019) implemented in Scanpy (Wolf et al. 2018). The number of selected genes is fixed at 2,000. We use the cluster assignment initialized as described above as the true labels and pre-train the annotator for 50 epochs.

Finally, we train the annotator together with the cluster assignment part (the encoder and decoder (e) & decoder(q)) for 25 epochs. In each epoch, the annotator is trained by using the clustering results as the true label for each cell. The output from the annotator $\mathbf{p}(\mathbf{r})$ is in turn used to evaluate the annotator loss $\mathcal{L}_{annotator}$ for the cluster assignment part. As described previously, the annotator is optimized using its own loss function, separately from the cluster assignment part.

There are two weighting hyperparameters in this model, $\gamma$ and $\tau$.

1. $\gamma$ which is used to weight the exclusive L1 penalty in $\mathcal{L}_{classification}$ of the annotator

   We employ a heuristic way to automatically select the value for $\gamma$ during the training process, without knowing the ground truth cell labels. During pre-training, the annotator will be easily overfitted if the number of features is as large or larger than the number of cells. This will be mitigated if we apply a more stringent regularization through a larger $\gamma$. Given a list of candidate values of $\gamma$, we start with a small value, pre-train the annotator and evaluate the classification accuracy on all available data (which we also use for training). If the accuracy is $\geq 0.99$, which indicates that the annotator is overfitted, we move on to the next larger value in the list and pre-train the annotator again until we find one resulting in the training accuracy $< 0.99$.

   The candidate values we consider range from 1e-3 to 2 when use both gene set activity scores and selected genes as input. When only gene set activity scores or genes are used as input, we start with 1e-4. As we show in Supplementary Results, our model is robust to different choices of starting values.

2. $\tau$ for the annotator loss in $\mathcal{L}_{\text{cluster}}$ in the cluster assignment part

   We set $\tau$ such that the magnitude of the annotator loss $\mathcal{L}_{annotator}$ is comparable to the magnitude of the reconstruction loss $\mathcal{L}_{\text{reconstruction}}$. We fix $\tau$ at 10 for all experiments. As we show in Supplementary Results, our model is robust to other choices of $\tau$.

##### 1.1.3 Details for Running the Prior Methods

For the prior methods, we use the default or recommending procedures / hyperparameters to all datasets unless we encounter difficuties in computaional resources or run time. For Leiden clustering, we use 30 neighbors and 32 dimensions for the principal components analysis (PCA). We fix the resolution at the default 1. For DESC, we adjust the hyperparameters for datasets with different sample sizes based on the recommending hyperparameters values listed in the Supplementary Table 2 in their manuscript (Li et al. 2020). We fix the resolution at 1 for the Louvain clustering used in DESC. For Seuratv3, we use the default top 2,000 most variable genes for the PCA. These genes were selected using Scanpy's implementation of Seuratv3-style variable genes selection (Wolf et al. 2018). For SIMLR, we use the default version and hyperparameter values for all datasets with a sample size $<$ 3,500. For datasets with larger sample size, we use their "large scale" version with the number of neighbors set at 30 and the number of dimensions for PCA set at 500. Given running their script to find the number of clusters for the larger datasets taking too much memory and cannot be finished, we set the number of clusters as the true number of cell types for the datasets with sample size $\geq$ 1e4. Running SIMLR for "pbmc68k" and "Atlas lung" takes too much memory and cannot be finished, even when we use 500 PCA as input instead of reading in the raw data. We thus don't have the results of SIMLR for "pbmc68k" and "Atlas lung". For SCCAF, we use 100 principal components learned from the preprocessed data, as mentioned in Miao et al. (2020). It is not clear from Miao et al. (2020) the specific value for k in k-nearest neighbors (k-NN) so we use the default value 15 from Wolf et al. (2018). Following Miao et al. (2020), we use Louvain (Blondel et al. 2008) clustering with resolution 1 to initialize the clusters. The other hyperparameter values are kept as default.

We use the default hyperparameter values for MARS and ItClust for all datasets. Given ItClust conducts gene filtration based on dispersion after taking into the input data, instead of the preprocessed data we use as input for other methods, we use raw count data without normalization for ItClust. For these raw count data, we filter the genes and cells under the same standards as for the preprocessed data to keep the cells and genes used as input for ItClust the same as for the others. We use stratified sampling to select cells for training for MARS and ItClust except for the liver and the lung dataset in the Tabula Muris where the least populated class has only one member and stratification is not allowed. For CellAssign, we use the implementation of it by scvi-tools (Gayoso et al. 2022). Following AW Zhang et al. (2019), we use raw count data as input. Following scvi-tools's tutorials, we set the size factor as the library size over the mean of library size and we also set a (pseudo-) cell type called "other" which does not have any of the marker genes expressed in the cell type marker gene table. We use markers from PanglaoDB (Franzén et al. 2019), where marker genes are grouped by tissues. We tried initially using all available cell type markers as input for any datasets and it turned out it runs extremely slow so we decided to use the ones from the tissue that best matches the tissue of the dataset. See Table S1 for details. It still takes a long time for running CellAssign on "pbmc68k" so we randomly sampled 70% of the cells stratified by true labels and run CellAssign on the sub-sampled data. We use all default hyperparameter values except for the batch size. We set the batch size as 1,024 (default) if the number of features after filtering is below 500, 512 if below 1,000 and 32 otherwise.

| dataset | matched tissue names in PanglaoDB |
|---|---|
| pbmc28k | Immune system |
| pbmc68k | Immune system |
| HuBMAP lymph_node | Immune system |
| HuBMAP spleen | Immune system |
| HuBMAP thymus | Immune system, Thymus |
| Atlas lung | Lungs |
| Tabula Muris Aorta | Vasculature |
| Tabula Muris Bladder | Urinary bladder |
| Tabula Muris Brain_Non-Myeloid | Brain |
| Tabula Muris Diaphragm | Epithelium |
| Tabula Muris Large_Intestine | GI tract |
| Tabula Muris Limb_Muscle | Skeletal muscle |
| Tabula Muris Tongue | Oral cavity |
| Tabula Muris Trachea | Vasculature |

Table S1: Dataset and their matched tissue names in PanglaoDB. The other tissues in the Tabula Muris dataset that are not shown here are matched with tissue names in PanglaoDB more straightforwardly (for example, Mammary_Gland to Mammary gland). We were unable to find matching tissue names for the adipose tissues in the Tabula Muris dataset so we did not run CellAssign for them.

###### 1.1.4 Details for Generating Random Values for "UNIFAN random"

For each feature, we randomly sample a value from $\mathcal{N}(0, 1)$. The total number of features equal to that of the default version ("UNIFAN genes & gene sets").

###### 1.1.5 Evaluation on the Enrichment of Cell Type Marker Sets in the Highly Weighted Genes

To interpret the highly weighted genes selected by the annotator, we select the genes whose coefficients above some thresholds. Given the cell type marker gene sets from MSigDB (Subramanian et al. 2005) (c8.all in MSigDB), we check the enrichment of these cell marker sets in the selected highly weighted genes by conducting binomial tests. For each cell marker gene set, we test if the chance of seeing its member appearing in the selected highly weighted genes is higher than the background rate, defined as the chance of seeing its member appearing in all genes used as features for the annotator. We use Benjamini/Hochberg procedure to correct for multiple testing and show the adjusted p-values in the results.

###### 1.1.6 Simulation Experiments

We conducted simulation experiments to test if UNIFAN is robust when cells contain novel pathways that have not been documented in the pathway database we use. We simulated expression data using pseudo-pathways that are not included in the pathway database. Specifically, we constructed pseudo-pathways by randomly selecting two pathways from the original database and combining them. We select from the pathways having number of genes larger or equal to 5 but smaller or equal to 20 (resulting in 5,036 pathways). Under this approach, we generated 3,000 pseudo-pathways. The number of genes per pseudo-pathway is about 20.

We then simulate expression data using these generated pseudo-pathways. For each simulation dataset, we generated 5 cell types, each having 300 cells. Each cell has 3,000 features (genes). For each cell type, we randomly selected a fixed number of pseudo-pathways. We combine the genes from these pseudo-pathways as the features. The rest of features are named as "PSEUDO1", "PSEUDO2", etc. We assume the expression of genes from a cell of a particular cell type following a multivariate normal distribution and set the mean values for the genes in the pseudo-pathways higher than the other genes (background genes). We set the mean for the background genes as 1. We assume all genes independent with each other and having variances equal to 1. We also applied a 10% dropout per cell to some of the simulation datasets.

We generated multiple simulation datasets under different number of selected pseudo-pathways (ranging from 15 to 25), different mean values for the pathway genes (2 or 3) and with or without drop-out. For each condition, we generated 5 replicates.

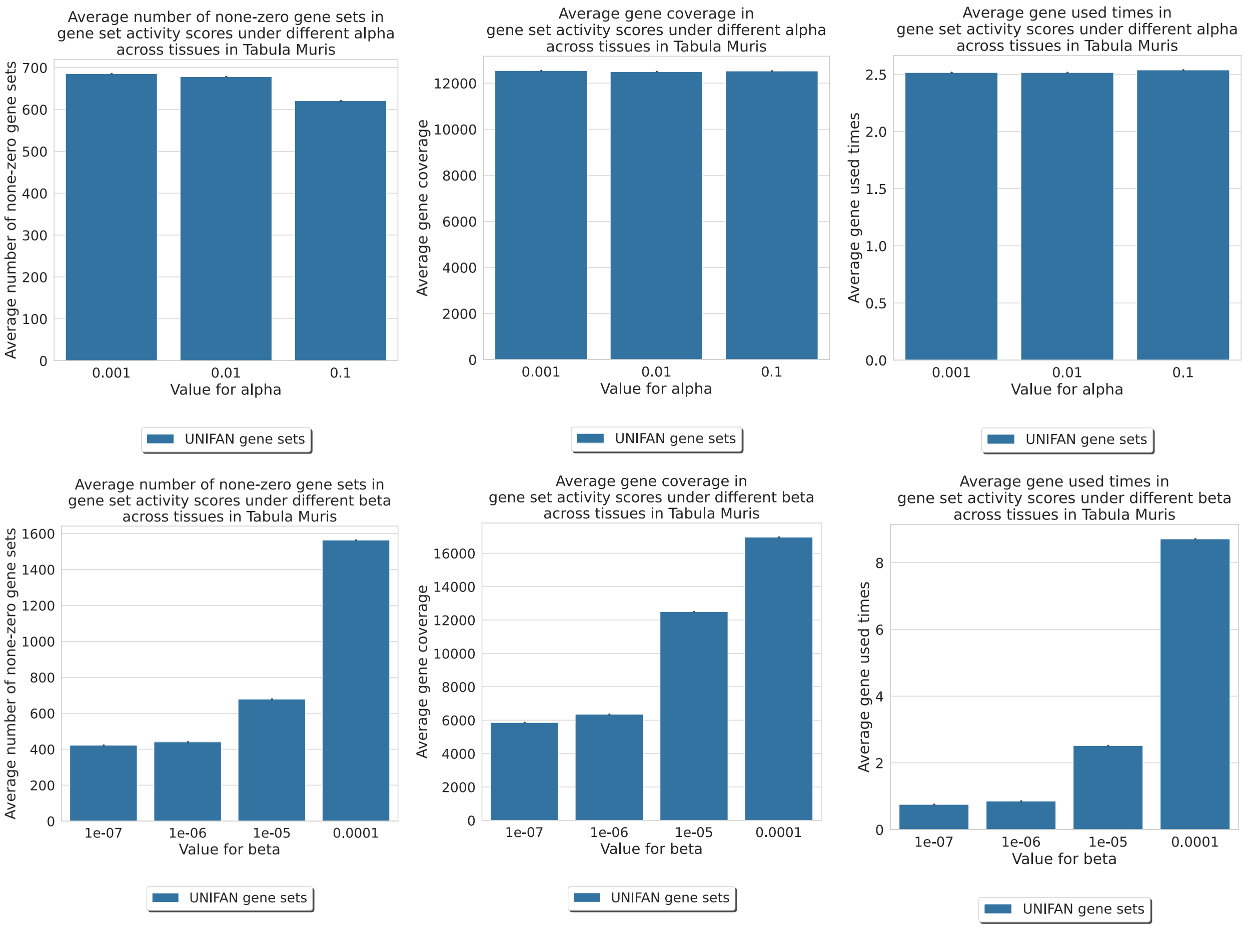

Figure S1: $\alpha$ and $\beta$'s impact on the gene set activity scores $\mathbf{r}$. For the values we tested, both $\alpha$ and $\beta$ impact on the sparsity of $\mathbf{r}$ (indicated by the number of non-zero gene sets in $\mathbf{r}$) while only $\beta$ impact on the gene coverage (indicated by the number of genes a cell used and the number of times a gene is used).

#### 1.2 Supplementary Results

##### 1.2.1 Alpha and Beta's Impact on Sparsity and Gene Coverage

##### 1.2.2 UNIFAN Robust to Different Choices of Hyperparameters

For the weighting hyperparameters $\alpha$, $\beta$, $\tau$ and $\gamma$, we vary the value for each of the hyperparameter at a time, while fixing the values for all other hyperparameters. We use all tissues in the Tabula Muris data for the evaluation. For each value, we run five times using different initializations. Given the values changing of $\alpha$ and $\beta$ may have different impact on the two variations of UNIFAN: "UNIFAN genes & gene sets" and "UNIFAN gene sets", we run both versions. For $\gamma$, given we employ the auto-selection of the value for $\gamma$ and we end up in the same $\gamma$ value regardless of the starting value for "UNIFAN genes & gene sets" and "UNIFAN genes", we only show the robust experiment results on "UNIFAN gene sets". For the others, we run the experiments on the default "UNIFAN genes & gene sets" version.

As shown in Figure S2, the clustering performance is similar under different hyperparameter choices within a set of reasonable values.

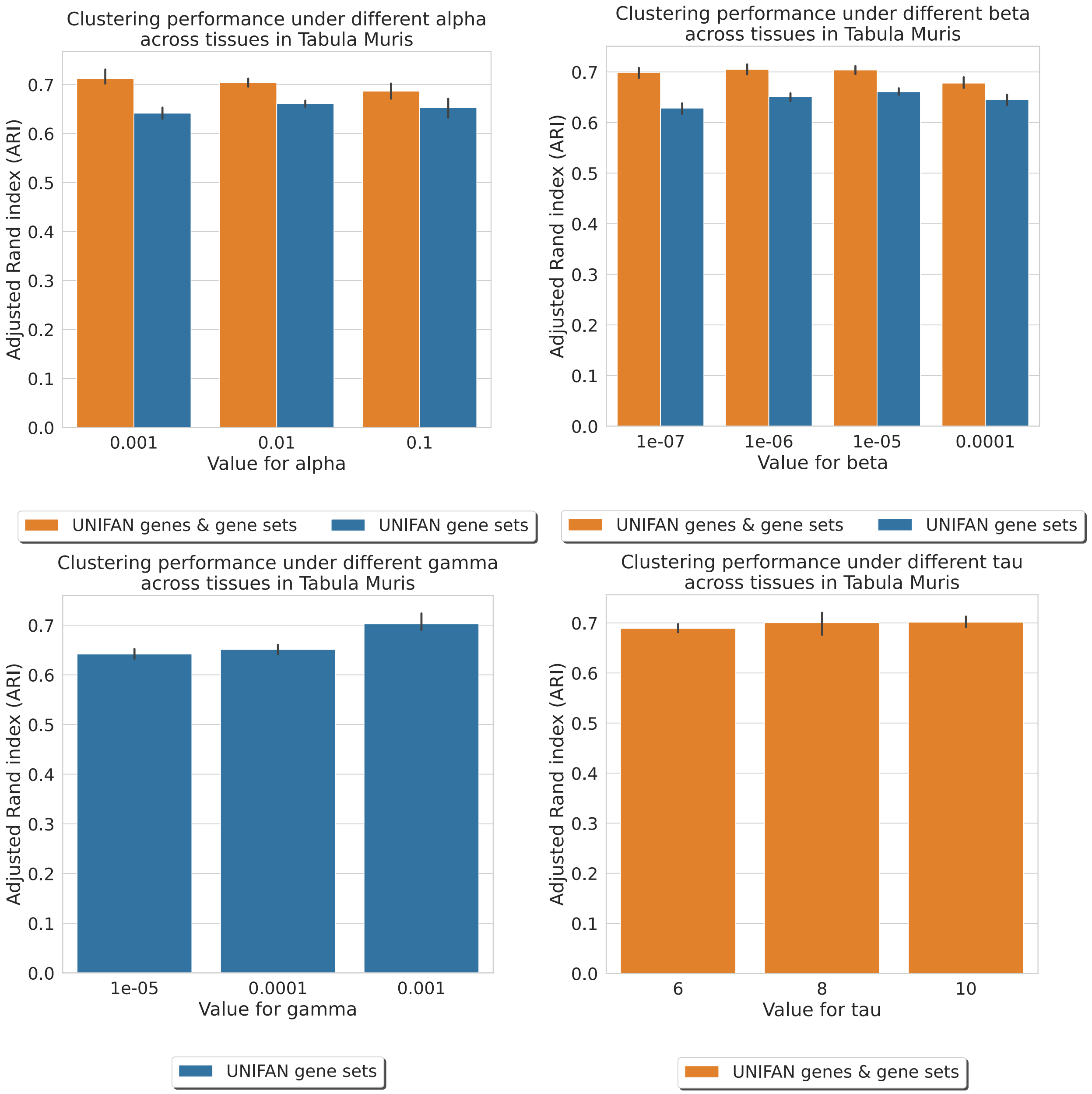

Figure S2: UNIFAN robust to different choices of hyperparameters. We varied the values for the weighting hyperparameters $\alpha$, $\beta$, $\tau$ and $\gamma$ and fixed all other hyperparameters. We run on all tissues in the Tabula Muris data for multiple times. The results indicate that our model is generally robust to different choices of the hyperparameters.

Other than the weighting hyperparameters, we also evaluate if our method is robust to different choices of the resolution value used in Leiden clustering initialization and to the number of epochs used for the joint training step of the clustering assignment part and the annotator. Again, we vary the value for each of these at a time, while fixing the values for the others. We use all tissues in the Tabula Muris data for the evaluation. For each value, we run five times using different initializations. We run the experiments on the default "UNIFAN genes & gene sets" version. We also check if the resulting number of clusters is affected by different choices of values.

As shown in Figure S3 and S4, both the resulting clustering performance and the number of clusters are similar under different choices of the resolution value and the number of epochs within a set of reasonable values.

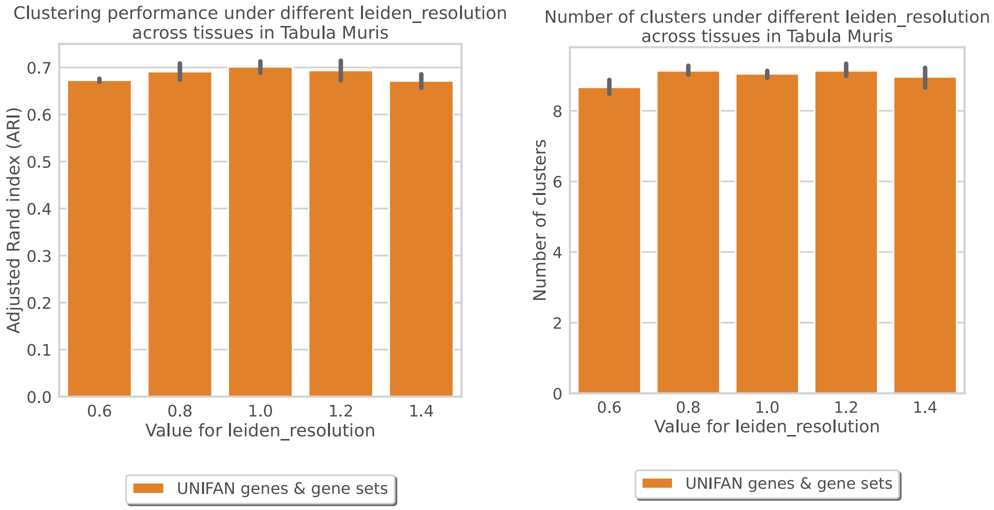

Figure S3: UNIFAN robust to different choices of the resolution value used in Leiden clustering initialization. We varied the values for the resolution and fixed all others. We run on all tissues in the Tabula Muris data for multiple times. The results indicate that our model is generally robust to different choices of the resolution value used in initialization.

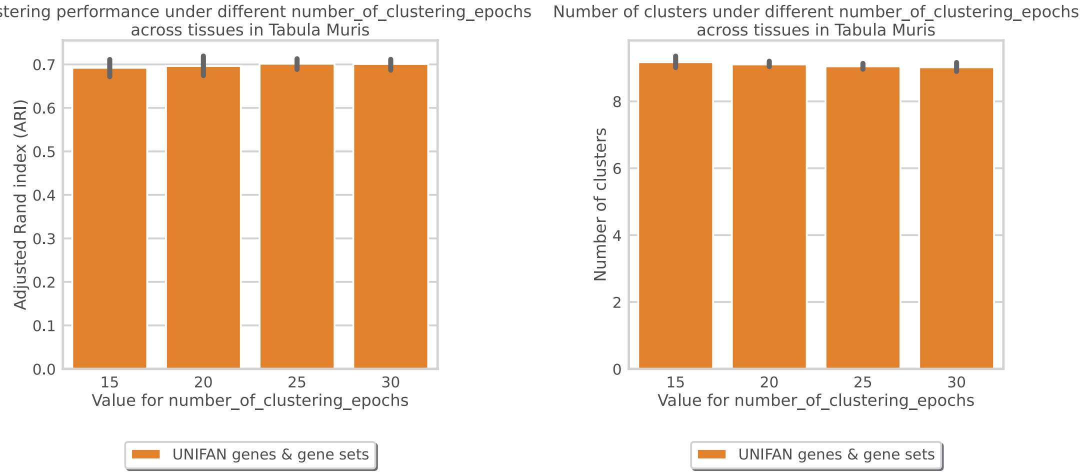

Figure S4: UNIFAN robust to different choices of the number of epochs for the joint training step. We varied the values for the training epochs and fixed all others. We run on all tissues in the Tabula Muris data for multiple times. The results indicate that our model is generally robust to different choices of the number of training epochs.

##### 1.2.3 Visualization and Interpretation of UNIFAN Results

| cluster | gene set name |
|---|---|
| 0 | GOBP ANTIGEN PROCESSING AND PRESENTATION ENDOGENOUS LIPID ANTIGEN VIA MHC CLASS IB |
| 3 | GOBP IMMUNE RESPONSE INHIBITING CELL SURFACE RECEPTOR SIGNALING PATHWAY |
| 3 | GOBP NEGATIVE REGULATION OF MACROPHAGE DERIVED FOAM CELL DIFFERENTIATION |
| 3 | GOBP REGULATION OF T CELL ACTIVATION VIA T CELL RECEPTOR CONTACT WITH ANTIGEN BOUND TO MHC MOLECULE ON ANTIGEN PRESENTING CELL |
| 4 | REACTOME ANTIGEN ACTIVATES B CELL RECEPTOR BCR LEADING TO GENERATION OF SECOND MESSENGERS |
| 5 | GOBP NEGATIVE REGULATION OF RESPIRATORY BURST INVOLVED IN INFLAMMATORY RESPONSE |
| 9 | GOBP POSITIVE REGULATION OF CD8 POSITIVE ALPHA BETA T CELL DIFFERENTIATION |

Table S2: Gene set names for highly weighted biological processes / pathways for “pbmc28k” that are truncated in Figure 2 D due to limit of space.

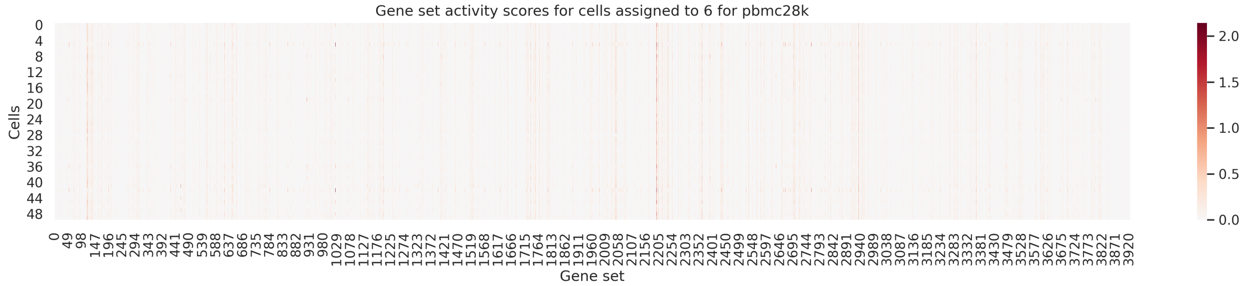

Figure S5: Other than interpreting from the annotator coefficients, we can also directly interpret the gene set activity scores to learn the biological processes or pathways important to cells / cell clusters. Here we show the gene set activity scores for cells assigned to cluster 6 in “pbmb28k” as an example. Each row corresponds to each cell’s gene set activity scores. We can clearly see that cells assigned to the same cluster tend to have similar gene set activity scores. The gene sets selected as important by the annotator also tend to have high scores. For example, most of the cells in cluster 6 are non-classical monocyte (ncMonocyte). The biological process directly relates to this cell type is “GOBP REGULATION OF INFLAMMATORY RESPONSE TO WOUNDING” (gene set at index 2298 in this plot), which is selected by the annotator and also among one of the highest scored gene sets according to the gene set activity scores. Note that due to space limit, here we only show the scores of 50 cells randomly selected from all cells assigned to cluster 6.

| cluster | gene set name |
|---|---|
| 0 | HAY BONE MARROW NAIVE T CELL |
| 0 | BUSSLINGER GASTRIC IMMUNE CELLS |
| 0 | FAN OVARY CL12 T LYMPHOCYTE NK CELL 2 |
| 0 | CUI DEVELOPING HEART C9 B T CELL |
| 0 | TRAVAGLINI LUNG CD4 MEMORY EFFECTOR T CELL |
| 0 | BUSSLINGER DUODENAL DIFFERENTIATING STEM CELLS |
| 1 | HAY BONE MARROW NK CELLS |
| 1 | TRAVAGLINI LUNG NATURAL KILLER CELL |
| 1 | RUBENSTEIN SKELETAL MUSCLE NK CELLS |
| 1 | DURANTE ADULT OLFACTORY NEUROEPITHELIUM NK CELLS |
| 1 | FAN EMBRYONIC CTX BRAIN EFFECTOR T CELL |
| 1 | AIZARANI LIVER C5 NK NKT CELLS 3 |
| 3 | TRAVAGLINI LUNG OLR1 CLASSICAL MONOCYTE CELL |
| 3 | TRAVAGLINI LUNG EREG DENDRITIC CELL |
| 3 | TRAVAGLINI LUNG CLASSICAL MONOCYTE CELL |
| 3 | HAY BONE MARROW IMMATURE NEUTROPHIL |
| 3 | FAN OVARY CL13 MONOCYTE MACROPHAGE |
| 3 | DURANTE ADULT OLFACTORY NEUROEPITHELIUM DENDRITIC CELLS |
| 4 | TRAVAGLINI LUNG B CELL |
| 4 | RUBENSTEIN SKELETAL MUSCLE B CELLS |
| 4 | DURANTE ADULT OLFACTORY NEUROEPITHELIUM B CELLS |
| 4 | FAN EMBRYONIC CTX BRAIN B CELL |
| 4 | AIZARANI LIVER C34 MHC II POS B CELLS |
| 4 | HAY BONE MARROW FOLLICULAR B CELL |
| 5 | AIZARANI LIVER C1 NK NKT CELLS 1 |
| 5 | FAN OVARY CL4 T LYMPHOCYTE NK CELL 1 |
| 5 | TRAVAGLINI LUNG CD8 NAIVE T CELL |
| 5 | BUSSLINGER GASTRIC IMMUNE CELLS |
| 5 | HAY BONE MARROW CD8 T CELL |
| 5 | DESCARTES FETAL PLACENTA LYMPHOID CELLS |
| 6 | HAY BONE MARROW MONOCYTE |
| 6 | TRAVAGLINI LUNG NONCLASSICAL MONOCYTE CELL |
| 6 | DESCARTES FETAL PANCREAS MYELOID CELLS |
| 6 | AIZARANI LIVER C6 KUPFFER CELLS 2 |
| 6 | DESCARTES FETAL CEREBELLUM MICROGLIA |
| 6 | AIZARANI LIVER C2 KUPFFER CELLS 1 |
| 7 | HAY BONE MARROW DENDRITIC CELL |
| 7 | BUSSLINGER ESOPHAGEAL DENDRITIC CELLS |
| 7 | DESCARTES FETAL INTESTINE MYELOID CELLS |
| 7 | FAN OVARY CL13 MONOCYTE MACROPHAGE |
| 7 | AIZARANI LIVER C2 KUPFFER CELLS 1 |
| 7 | AIZARANI LIVER C25 KUPFFER CELLS 4 |
| 9 | HAY BONE MARROW DENDRITIC CELL |
| 9 | TRAVAGLINI LUNG PLASMACYTOID DENDRITIC CELL |
| 9 | FAN OVARY CL18 B LYMPHOCYTE |

Table S3: Gene set names for the enriched cell type marker sets in the highly weighted genes for “pbmc28k”. These names are truncated in Figure 2 E due to limit of space.

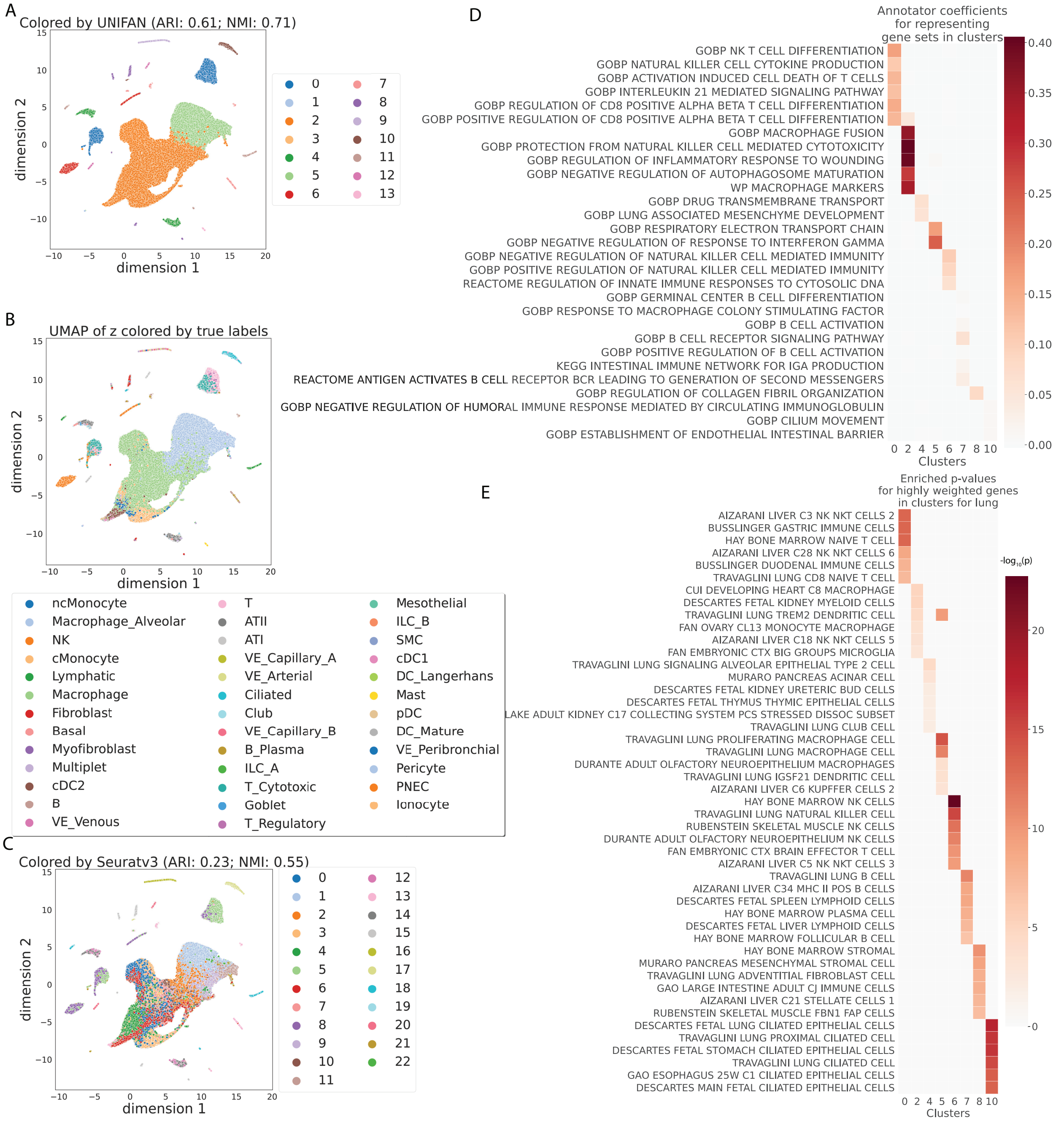

Figure S6: UNIFAN accurately clusters cells and correctly identifies biological processes / pathways. Results presented for the "Atlas lung" dataset. **A, B and C**: UMAP visualization of the low-dimensional representation $\mathbf{z_e}$ of cells output from UNIFAN. **A**: Colored by the clusters found by UNIFAN; **B**: colored by the true cell type labels; **C**: colored by the clusters found by Seuratv3. **D**: Coefficients learned by the annotator for highly weighted gene sets for some of the clusters. **E**: Enrichment p-values of cell type marker sets in the highly weighted genes learned by the annotator. Here we show the result from the best run out of multiple runs for UNIFAN and for Seuratv3.

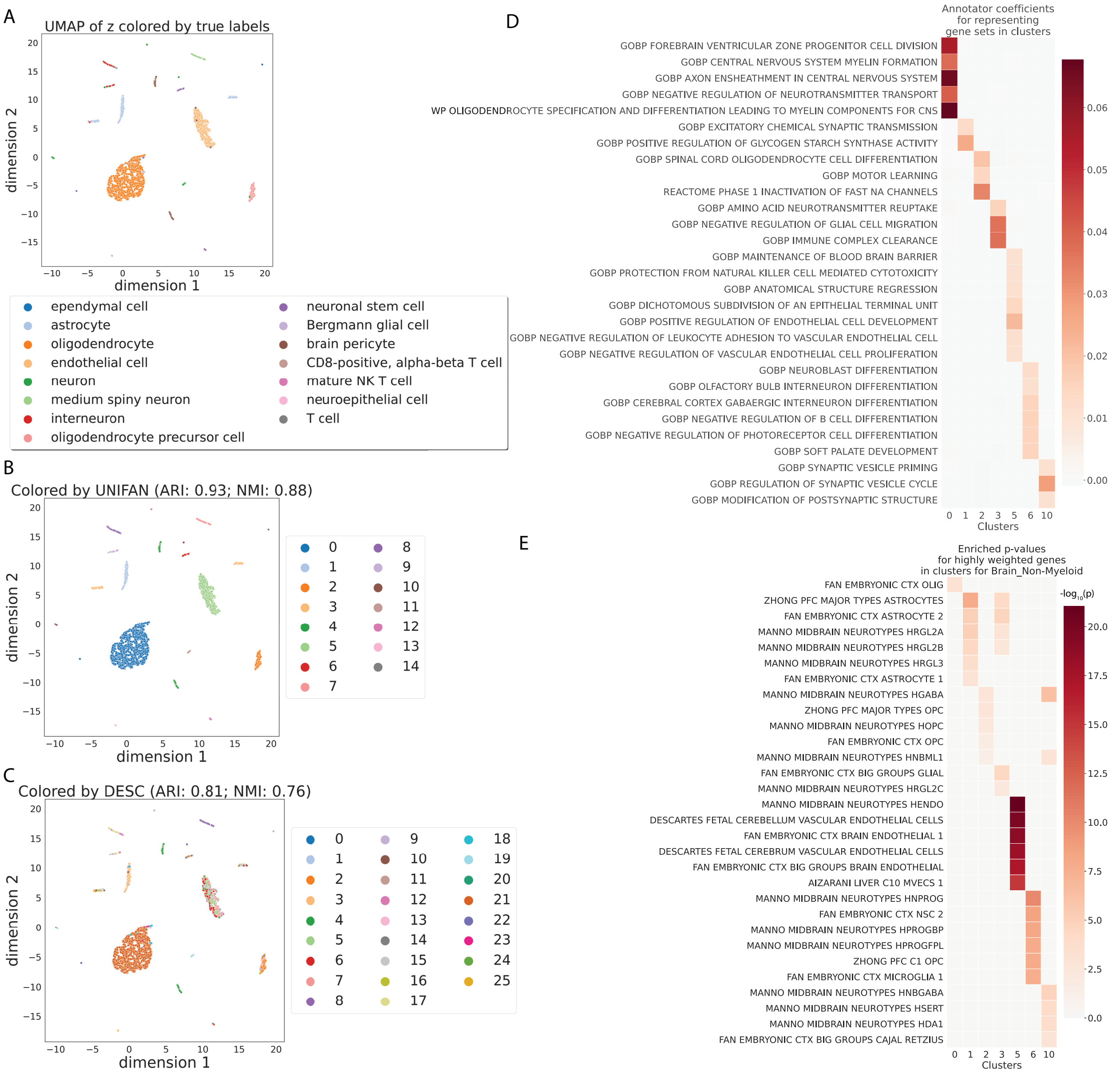

Figure S7: UNIFAN accurately clusters cells and correctly identifies biological processes / pathways. Results presented for the "Tabula Muris Brain_Non-Myeloid" dataset. **A, B and C**: UMAP visualization of the low-dimensional representation $\mathbf{z_e}$ of cells output from UNIFAN. **A**: Colored by the true cell type labels; **B**: colored by the clusters found by UNIFAN; **C**: colored by the clusters found by DESC. **D**: Coefficients learned by the annotator for highly weighted gene sets for some of the clusters. **E**: Enrichment p-values of cell type marker sets in the highly weighted genes learned by the annotator. Here we show the result from the best run out of multiple runs for UNIFAN and for DESC.

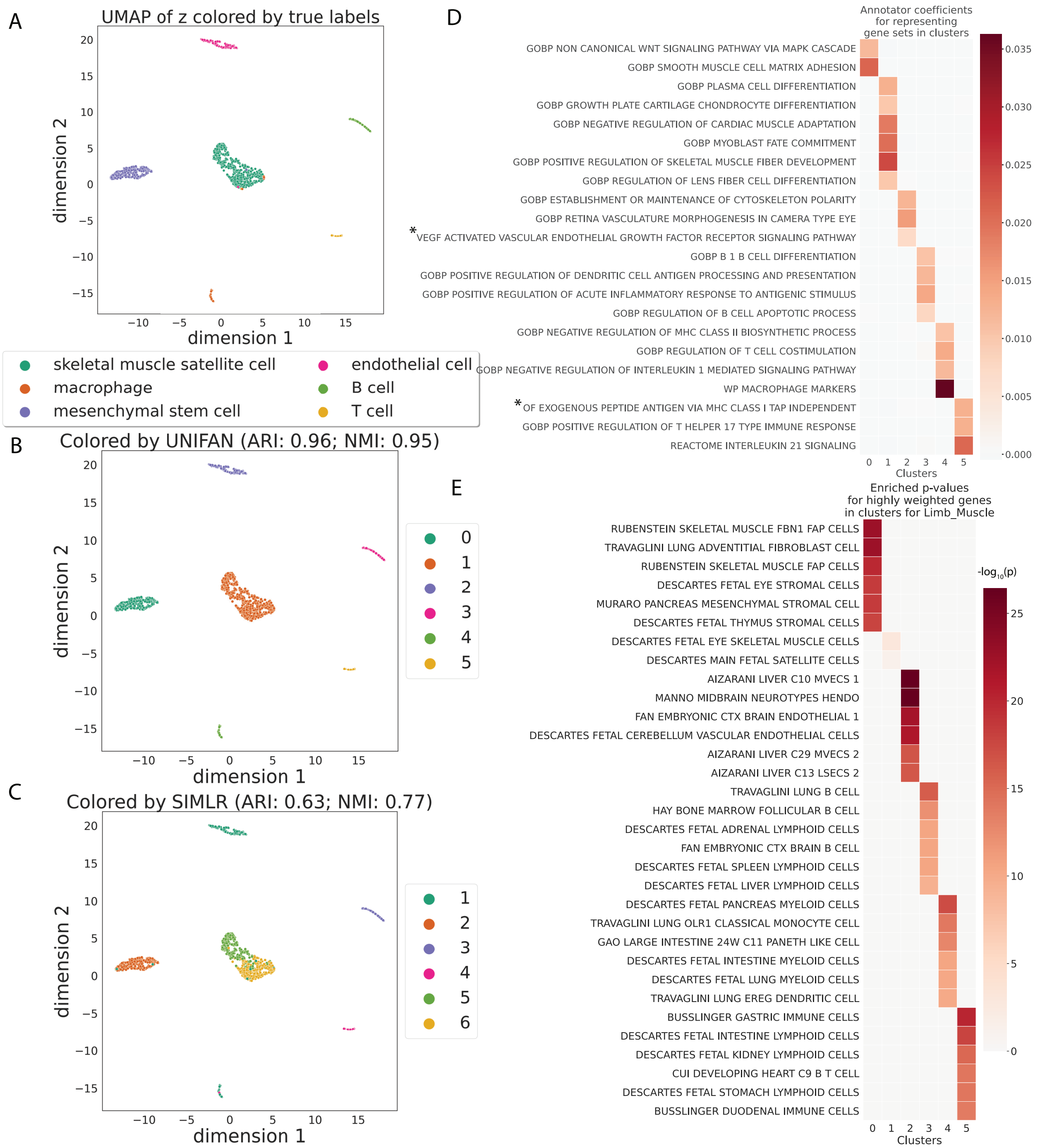

Figure S8: UNIFAN accurately clusters cells and correctly identifies biological processes / pathways. Results presented for the "Tabula Muris Limb_Muscle" dataset. **A, B and C**: UMAP visualization of the low-dimensional representation $\mathbf{z_e}$ of cells output from UNIFAN. **A**: Colored by the true cell type labels; **B**: colored by the clusters found by UNIFAN; **C**: colored by the clusters found by SIMLR. **D**: Coefficients learned by the annotator for highly weighted gene sets for some of the clusters. **E**: Enrichment p-values of cell type marker sets in the highly weighted genes learned by the annotator. Here we show the result from the best run out of multiple runs for UNIFAN and for SIMLR. The truncated gene sets (marked by *) are: "GOBP POSITIVE REGULATION OF ENDOTHELIAL CELL CHEMOTAXIS BY VEGF ACTIVATED VASCULAR ENDOTHELIAL GROWTH FACTOR RECEPTOR SIGNALING PATHWAY" and "GOBP ANTIGEN PROCESSING AND PRESENTATION OF EXOGENOUS PEPTIDE ANTIGEN VIA MHC CLASS I TAP INDEPENDENT".

##### 1.2.4 Compare UNIFAN Variations and UNIFAN with Prior Methods

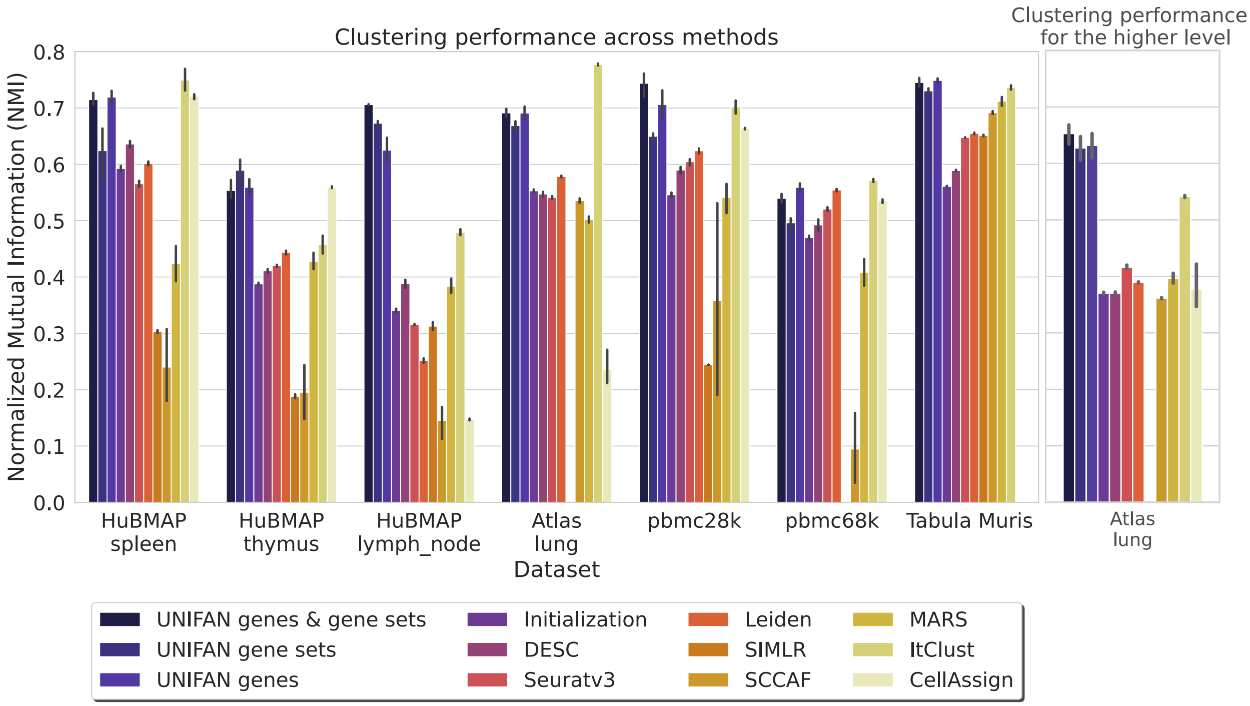

Figure S9: UNIFAN significantly outperforms other methods. "UNIFAN genes & gene sets" is the default UNIFAN version using both gene set activity scores and a subset of genes as features for the annotator; "UNIFAN gene sets" and "UNIFAN genes" uses only the gene set activity scores and the gene subset respectively. "Initialization" is the initialization clustering results. The others are the prior methods we used for comparison. For the Tabula Muris data, we take the average over all tissues. See Figure S13 and S14 for the per tissue results. The "Atlas lung" data provides two levels of cell type annotations and so we show results for both (less detailed annotation comparison shown on the right). SIMLR was unable to cluster the "pbmc68k" and "Atlas lung" data since it ran out of memory. CellAssign does not have an average over all tissues for "Tabula Muris" because it was unable to annotate some tissues which do not have matched cell type marker genes in the database (e.g., adipose tissues). See Supplementary Methods for details.

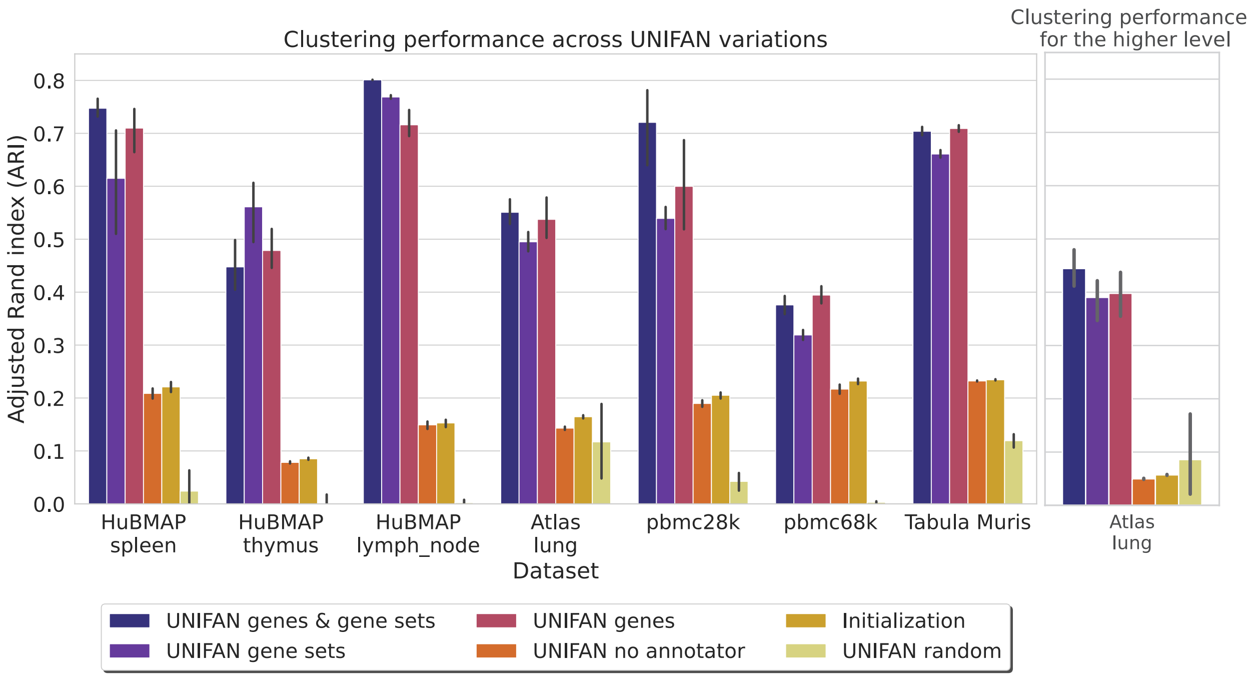

Figure S10: "UNIFAN genes & gene sets" (default version) performs the best most of the time and both the annotator and its features (gene set activity scores and genes) are crucial to UNIFAN's performance. "UNIFAN genes & gene sets" uses both gene set activity scores and a subset of genes as features for the annotator; "UNIFAN gene sets" and "UNIFAN genes" uses only the gene set activity scores and the gene subset respectively. "UNIFAN no annotator" does not use the annotator ($\tau = 0$). "UNIFAN random" use randomly generated values as features for the annotator (the dimension of these values is the same as the default version). "Initialization" is the initialization clustering results. For the Tabula Muris data, we take the average over all tissues. The "Atlas lung" data provides two levels of cell type annotations and so we show results for both (less detailed annotation comparison shown on the right).

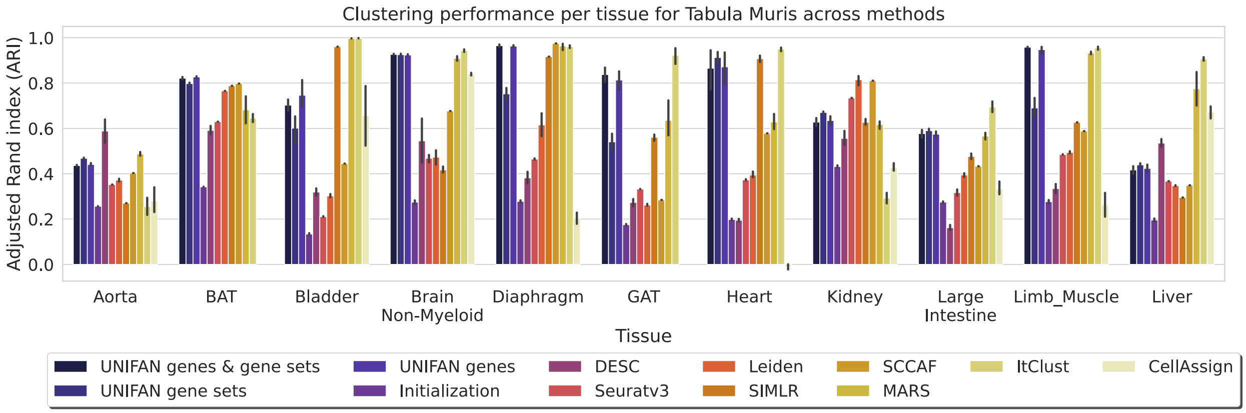

Figure S11: Compare per tissue result for Tabula Muris among methods using ARI - the first 11 tissues. "UNIFAN genes & gene sets" is the default UNIFAN version using both gene set activity scores and a subset of genes as features for the annotator; "UNIFAN gene sets" and "UNIFAN genes" uses only the gene set activity scores and the gene subset respectively. "Initialization" is the initialization clustering results. The others are the prior methods we used for comparison.

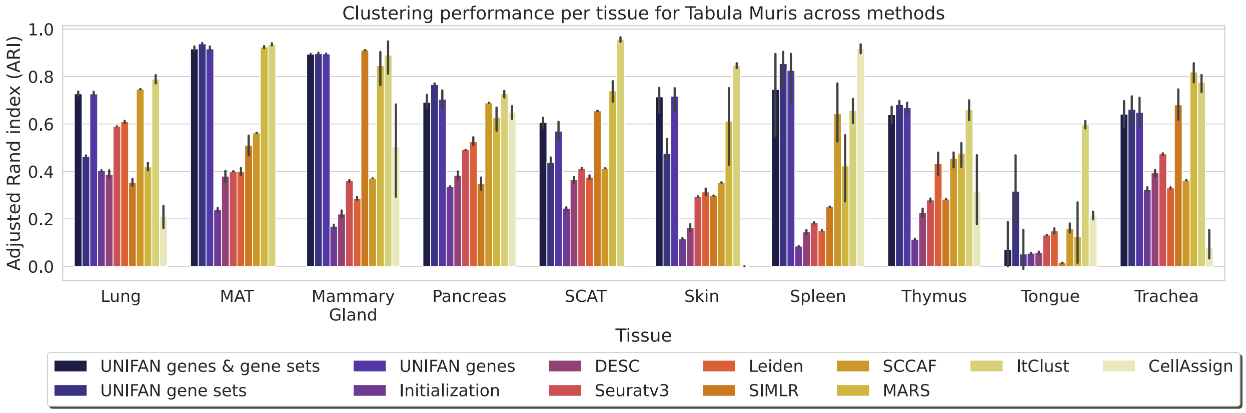

Figure S12: Compare per tissue result for Tabula Muris among methods using ARI - the rest 10 tissues. "UNIFAN genes & gene sets" is the default UNIFAN version using both gene set activity scores and a subset of genes as features for the annotator; "UNIFAN gene sets" and "UNIFAN genes" uses only the gene set activity scores and the gene subset respectively. "Initialization" is the initialization clustering results. The others are the prior methods we used for comparison.

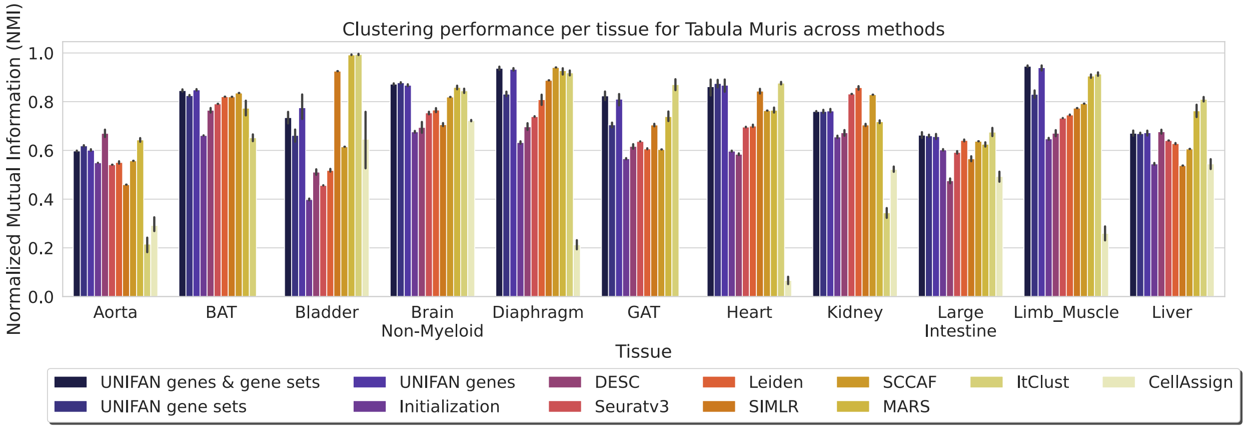

Figure S13: Compare per tissue result for Tabula Muris among methods using NMI - the first 11 tissues. "UNIFAN genes & gene sets" is the default UNIFAN version using both gene set activity scores and a subset of genes as features for the annotator; "UNIFAN gene sets" and "UNIFAN genes" uses only the gene set activity scores and the gene subset respectively. "Initialization" is the initialization clustering results. The others are the prior methods we used for comparison.

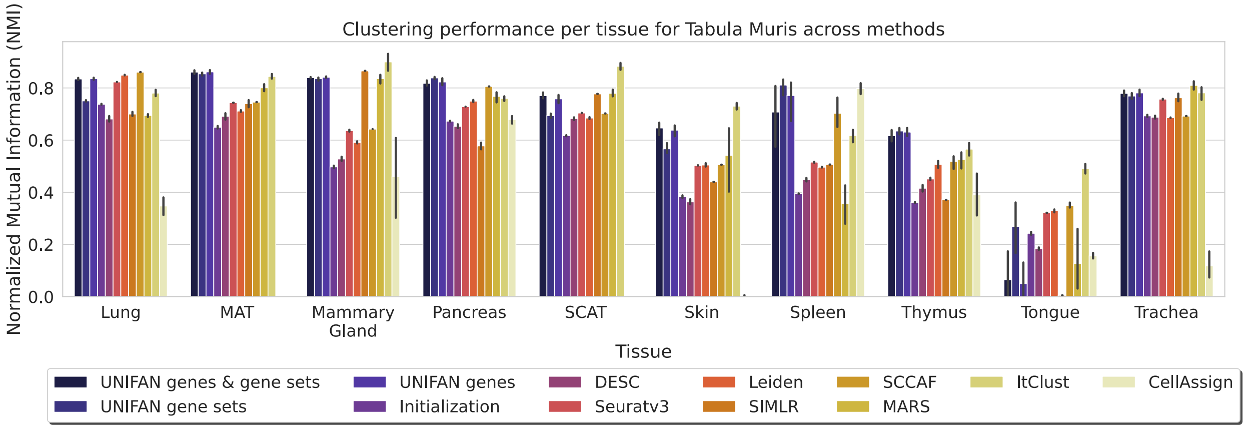

Figure S14: Compare per tissue result for Tabula Muris among methods using NMI - the rest 10 tissues. "UNIFAN genes & gene sets" is the default UNIFAN version using both gene set activity scores and a subset of genes as features for the annotator; "UNIFAN gene sets" and "UNIFAN genes" uses only the gene set activity scores and the gene subset respectively. "Initialization" is the initialization clustering results. The others are the prior methods we used for comparison.

##### 1.2.5 Models are Transferable Across Tissues and Species

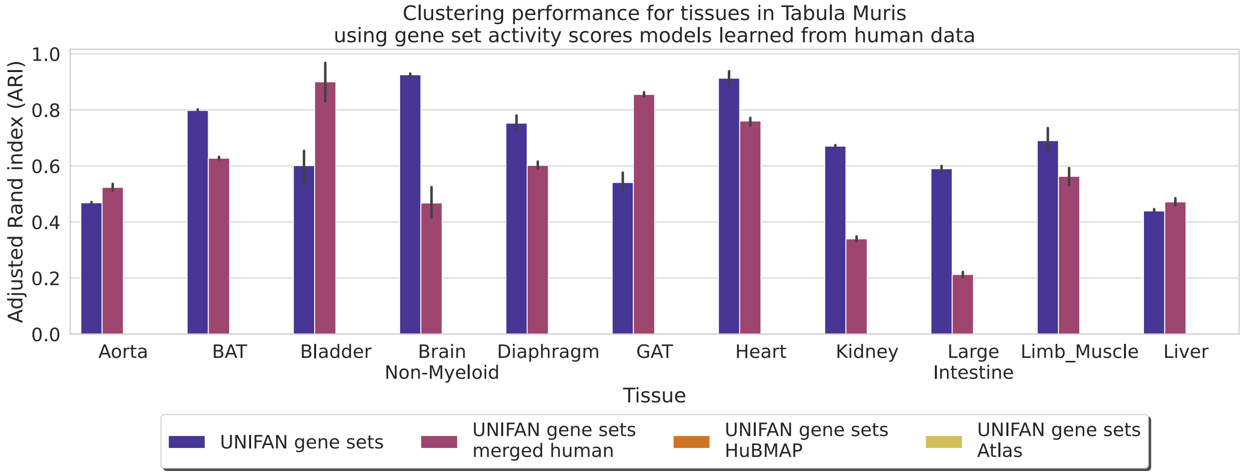

Figure S15: Compare per tissue result for Tabula Muris using different gene set activity scores models - the first 11 tissues. All versions of UNIFAN methods use models pre-trained on human tissues except for "UNIFAN gene sets", which used models trained on the same datasets as we discussed before. "UNIFAN gene sets merged human" uses the model pre-trained on all available human tissues. "UNIFAN gene sets HuBMAP" uses the model pre-trained on the corresponding HuBMAP tissue (HuBMAP spleen or thymus). "UNIFAN gene sets Atlas" uses the model pre-trained on the "Atlas lung" dataset.

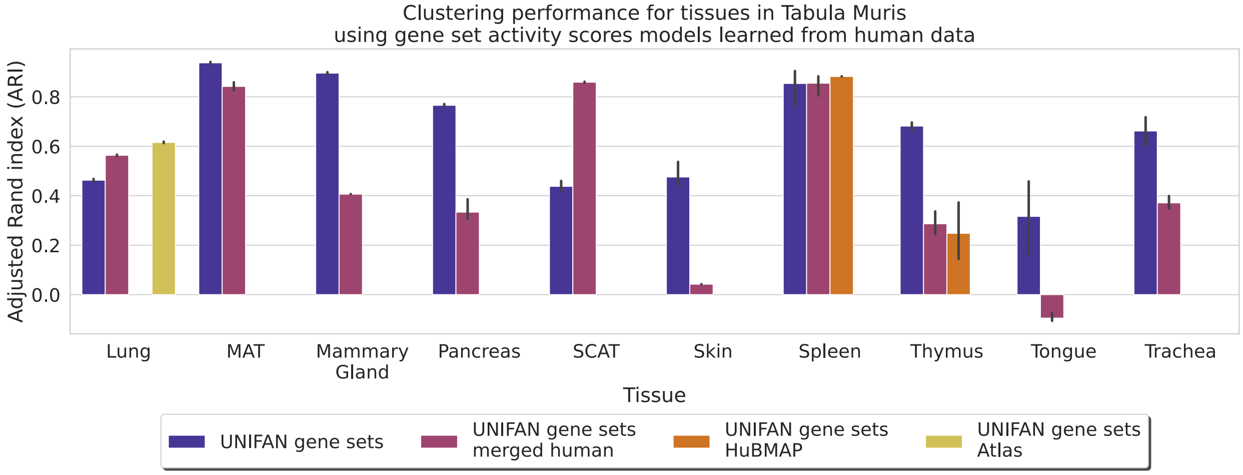

Figure S16: Compare per tissue result for Tabula Muris using different gene set activity scores models - the rest 10 tissues. All versions of UNIFAN methods use models pre-trained on human tissues except for "UNIFAN gene sets", which used models trained on the same datasets as we discussed before. "UNIFAN gene sets merged human" uses the model pre-trained on all available human tissues. "UNIFAN gene sets HuBMAP" uses the model pre-trained on the corresponding HuBMAP tissue (HuBMAP spleen or thymus). "UNIFAN gene sets Atlas" uses the model pre-trained on the "Atlas lung" dataset.

###### 1.2.6 UNIFAN Robust to Technical Variations

While we did not purposely design our method to tackle technical variations such as batch effect, we found UNIFAN to be robust to non-biological variations in many cases. Figure S17 shows the visualization and clustering results of the "HuBMAP lymph_node" dataset. Figure S17A shows the UMAP visualization of the initial low-dimensional representations of the cells output from a standard autoencoder. We see cells labeled by the same cell types segregated in the low-dimensional space. While other clustering methods such as DESC are impacted by this and failed to cluster cells from the same type together (Figure S17B), UNIFAN successfully clustered them together as shown by the UMAP visualization of the low-dimensional representations learned by UNIFAN (Figure S17D and S17E). This may be attributed to our way of using the gene set activity scores to guide the clustering decisions, which allows UNIFAN to focus on more relevant co-expression of genes and overcome noise attributed to technical variations.

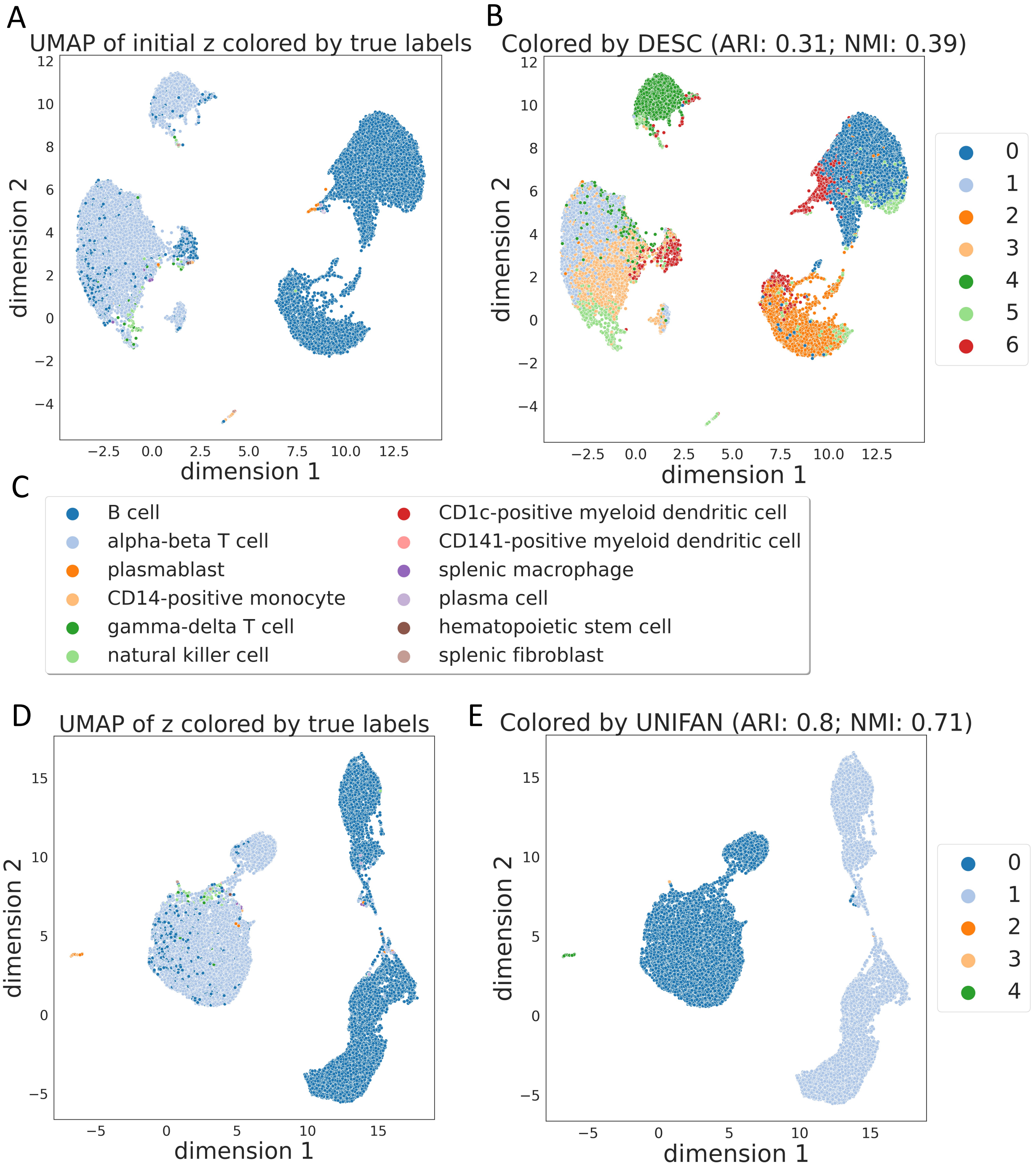

Figure S17: UNIFAN overcomes technical variations when clustering "HuBMAP lymph_node". **A, B**: UMAP visualization of the initial low-dimensional representation of cells output from a standard autoencoder; **A**: Colored by true cell type labels; **B**: colored by results from DESC. **C**: Legend for the visualization plots **A, D** colored by the true labels. **D, E**: UMAP visualization of the low-dimensional representation $\mathbf{z_e}$ of cells output from UNIFAN; **D**: Colored by true cell type labels; **E**: colored by the clusters found by UNIFAN.

##### 1.2.7 UNIFAN Robust to Novel Cell Types with Novel Pathways

We conducted simulation experiments to test if UNIFAN is robust when cells contain novel pathways that have not been documented in the pathway database we use. We applied UNIFAN to the simulation datasets generated as described in section 1.1.6. Given we mainly focus on robustness on novel pathways in this simulation experiment, we only ran the UNIFAN version which uses only gene set activity scores as features for the annotator ("UNIFAN gene sets"). To evaluate UNIFAN's performance, we consider two metrics. The first one is adjusted Rand index (ARI) to assess clustering performance. The second one is the ratio of pathways we used that are highly-weighted by UNIFAN over all (real) pathways we used to generate a cell type. Specifically, given we construct pseudo-pathways by combining two randomly selected real pathways, we break the pseudo-pathways we used to generated a cell type into two and check how many of these are overlapped with the highly-weighted gene sets found by UNIFAN. By highly-weighted, we mean those having coefficients larger than 90% quantile of all coefficients for the corresponding cluster.

As Figure S18 shows, our method is generally robust to the cases where cell types contain pathways that are not included in the current database. We observed that under different conditions (e.g., different number of selected pseudo-pathways), the clusters we obtained always correspond to the true cell types well (the adjusted Rand index is almost 1). Among all (real) pathways we used to generate a cell type, about 30% of them are ranked high by UNIFAN.

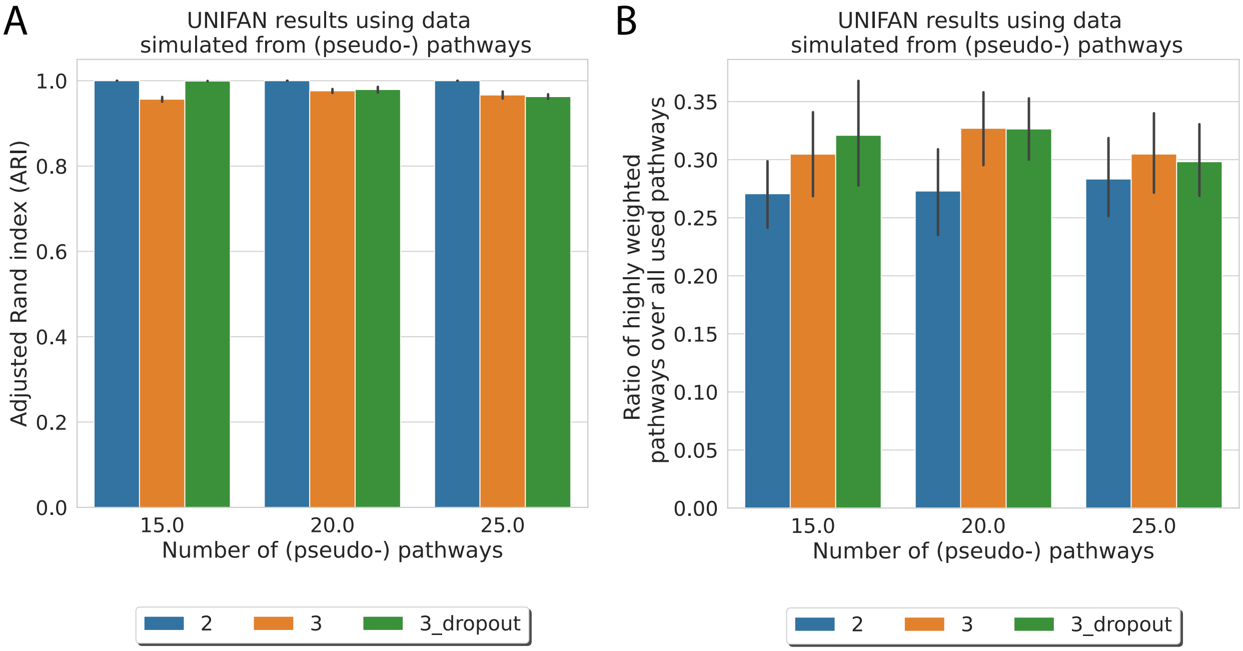

Figure S18: Simulation results on UNIFAN's robustness to novel cell types with novel pathways. **A**: adjusted Rand index (ARI) of UNIFAN's results on the simulation data; **B**: percent of (real) pathways used to generate a cell type ranked high by UNIFAN. The values are averaged over all replicates and all cell types. 2-when using 2 as the mean expression values for genes in pseudo-pathways; 3-when using 3 as the mean expression values for genes in pseudo-pathways; 3_dropout-when using 3 as the mean expression values for genes in pseudo-pathways and applying 10% dropout;

##### 1.2.8 Runtime

We recorded the runtime of UNIFAN for several representative datasets and the results are shown in Table S4. We also recorded the runtime of an unsupervised scRNA-Seq clustering method DESC (Li et al. 2020) and a cell type assignment method based on known markers CellAssign (AW Zhang et al. 2019), as comparisons. All results reported in this section are from experiments conducted in Linux Mint 19 with Intel(R) Xeon(R) W-2123 CPU @ 3.60GHz and 16 GB memory without using any GPU.

| Dataset | Number of cells | Number of genes | Time for training gene set activity scores model | Time for clustering | Time for DESC | Time for CellAssign |
|---|---|---|---|---|---|---|
| Tabula Muris Aorta | 366 | 22,904 | 13 min | 13 min | 26 s | 1 min 38 s |
| Tabula Muris SCAT | 1,721 | 22,904 | 51 min | 17 min | 2 min 18 s | - |
| Tabula Muris Heart | 4,433 | 22,904 | 2 hrs 8 min | 30 min | 5 min 54 s | 5 min 10 s |
| pbmc28k | 25,185 | 19,404 | 2 hrs 51 min | 2 hrs 30 min | 28 min | 14 hrs 1 min |
| Atlas lung | 96,282 | 17,315 | 10 hrs 17 min | 4 hrs 30 min | 1 hr 26 min | 3 hrs 43 min |

Table S4: UNIFAN, DESC and CellAssign runtime for datasets with various sample and feature sizes. For UNIFAN, the runtime is recorded in the two separate steps - training the gene set activity scores model and clustering. The training process for the gene set activity scores model consumes most of the time. We were unable to run CellAssign on "Tabula Muris SCAT" because this tissue does not have matched cell type marker genes in the marker database.

#### References

Abdelaal T, Michielsen L, Cats D, Hoogduin D, Mei H, Reinders MJ, and Mahfouz A. 2019. A comparison of automatic cell identification methods for single-cell RNA sequencing data. *Genome Biol.* **20**: 1–19.

Adams TS et al. 2020. Single-cell RNA-seq reveals ectopic and aberrant lung-resident cell populations in idiopathic pulmonary fibrosis. *Sci. Adv.* **6**: eaba1983.

Alavi A, Ruffalo M, Parvangada A, Huang Z, and Bar-Joseph Z. 2018. A web server for comparative analysis of single-cell RNA-seq data. *Nat. Commun.* **9**: 1–11.

Becht E, McInnes L, Healy J, Dutertre CA, Kwok IW, Ng LG, Ginhoux F, and Newell EW. 2019. Dimensionality reduction for visualizing single-cell data using UMAP. *Nat. Biotechnol.* **37**: 38–44.

Blondel VD, Guillaume JL, Lambiotte R, and Lefebvre E. 2008. Fast unfolding of communities in large networks. *J. Stat. Mech: Theory Exp.* **2008**: P10008.

Brbić M, Zitnik M, Wang S, Pisco AO, Altman RB, Darmanis S, and Leskovec J. 2020. MARS: discovering novel cell types across heterogeneous single-cell experiments. *Nat. Methods.* **17**: 1200–1206.

Clarke ZA, Andrews TS, Atif J, Pouyabahar D, Innes BT, MacParland SA, and Bader GD. 2021. Tutorial: guidelines for annotating single-cell transcriptomic maps using automated and manual methods. *Nat. Protoc.* **16**: 2749–2764.

Consortium H et al. 2019. The human body at cellular resolution: the NIH Human Biomolecular Atlas Program. *Nature.* **574**: 187.

Consortium TM et al. 2018. Single-cell transcriptomics of 20 mouse organs creates a Tabula Muris. *Nature.* **562**: 367–372.

Ernst J, Vainas O, Harbison CT, Simon I, and Bar-Joseph Z. 2007. Reconstructing dynamic regulatory maps. *Mol. Syst. Biol.* **3**: 74.

Fields RD. 2008. Oligodendrocytes changing the rules: action potentials in glia and oligodendrocytes controlling action potentials. *The Neuroscientist.* **14**: 540–543.

Fortuin V, Hüser M, Locatello F, Strathmann H, and Rätsch G. 2018. Som-vae: Interpretable discrete representation learning on time series. *arXiv preprint arXiv:1806.02199.*

Franzén O, Gan LM, and Björkegren JL. 2019. PanglaoDB: a web server for exploration of mouse and human single-cell RNA sequencing data. *Database.* **2019**:

Gayoso A, Lopez R, Xing G, Boyeau P, Valiollah Pour Amiri V, Hong J, Wu K, Jayasuriya M, Mehlman E, Langevin M, et al. 2022. A Python library for probabilistic analysis of single-cell omics data. *Nat. Biotechnol.* 1–4.

Hu J, Li X, Hu G, Lyu Y, Susztak K, and Li M. 2020. Iterative transfer learning with neural network for clustering and cell type classification in single-cell RNA-seq analysis. *Nat. Mach. Intell.* **2**: 607–618.

Kingma DP and Ba J 2014. Adam: A Method for Stochastic Optimization. eprint: `arXiv:1412.6980`.

Kiselev VY, Yiu A, and Hemberg M. 2018. scmap: projection of single-cell RNA-seq data across data sets. *Nat. Methods.* **15**: 359–362.

Li X, Wang K, Lyu Y, Pan H, Zhang J, Stambolian D, Susztak K, Reilly MP, Hu G, and Li M. 2020. Deep learning enables accurate clustering with batch effect removal in single-cell RNA-seq analysis. *Nat. Commun.* **11**: 1–14.

Lu Y, Rosenfeld R, Simon I, Nau GJ, and Bar-Joseph Z. 2008. A probabilistic generative model for GO enrichment analysis. *Nucleic Acids Res.* **36**: e109–e109.

Miao Z, Moreno P, Huang N, Papatheodorou I, Brazma A, and Teichmann SA. 2020. Putative cell type discovery from single-cell gene expression data. *Nat. Methods.* **17**: 621–628.

Oord Avd, Vinyals O, and Kavukcuoglu K. 2017. Neural discrete representation learning. *arXiv preprint arXiv:1711.00937*.

Paszke A, Gross S, Chintala S, Chanan G, Yang E, DeVito Z, Lin Z, Desmaison A, Antiga L, and Lerer A. 2017. Automatic differentiation in PyTorch.

Pedregosa F et al. 2011. Scikit-learn: Machine Learning in Python. *J. Mach. Learn. Res.* **12**: 2825–2830.

Pliner HA, Shendure J, and Trapnell C. 2019. Supervised classification enables rapid annotation of cell atlases. *Nat. Methods.* **16**: 983–986.

Schmidl C, Renner K, Peter K, Eder R, Lassmann T, Balwierz PJ, Itoh M, Nagao-Sato S, Kawaji H, Carninci P, et al. 2014. Transcription and enhancer profiling in human monocyte subsets. *Blood.* **123**: e90–e99.

Stuart T, Butler A, Hoffman P, Hafemeister C, Papalexi E, Mauck III WM, Hao Y, Stoeckius M, Smibert P, and Satija R. 2019. Comprehensive integration of single-cell data. *Cell.* **177**: 1888–1902.

Subramanian A et al. 2005. Gene set enrichment analysis: A knowledge-based approach for interpreting genome-wide expression profiles. *Proc. Natl. Acad. Sci. U. S. A.* **102**: 15545–15550.

Tanay A and Regev A. 2017. Scaling single-cell genomics from phenomenology to mechanism. *Nature.* **541**: 331–338.

Traag VA, Waltman L, and Van Eck NJ. 2019. From Louvain to Leiden: guaranteeing well-connected communities. *Sci. Rep.* **9**: 1–12.

van der Maaten L and Hinton G. 2008. Visualizing High-Dimensional Data Using t-SNE. *J. Mach. Learn. Res.* **9**: 2579–2605.

Van Der Wijst MG, Brugge H, De Vries DH, Deelen P, Swertz MA, and Franke L. 2018. Single-cell RNA sequencing identifies celltype-specific cis-eQTLs and co-expression QTLs. *Nat. Genet.* **50**: 493–497.

Wang B, Zhu J, Pierson E, Ramazzotti D, and Batzoglou S. 2017. Visualization and analysis of single-cell RNA-seq data by kernel-based similarity learning. *Nat. Methods.* **14**: 414–416.

Wei Z and Zhang S. 2021. CALLR: a semi-supervised cell-type annotation method for single-cell RNA sequencing data. *Bioinformatics.* **37**: i51–i58.

Wolf FA, Angerer P, and Theis FJ. 2018. SCANPY: large-scale single-cell gene expression data analysis. *Genome Biol.* **19**: 1–5.

Wosczyna MN, Konishi CT, Carbajal EEP, Wang TT, Walsh RA, Gan Q, Wagner MW, and Rando TA. 2019. Mesenchymal stromal cells are required for regeneration and homeostatic maintenance of skeletal muscle. *Cell Rep.* **27**: 2029–2035.

Xie J, Girshick R, and Farhadi A 2016. Unsupervised deep embedding for clustering analysis. In: *International conference on machine learning.* PMLR, pp. 478–487.

Xin J, Mark A, Afrasiabi C, Tsueng G, Juchler M, Gopal N, Stupp GS, Putman TE, Ainscough BJ, Griffith OL, et al. 2016. High-performance web services for querying gene and variant annotation. *Genome Biol.* **17**: 1–7.

Zhang AW, O'Flanagan C, Chavez EA, Lim JL, Ceglia N, McPherson A, Wiens M, Walters P, Chan T, Hewitson B, et al. 2019. Probabilistic cell-type assignment of single-cell RNA-seq for tumor microenvironment profiling. *Nat. Methods.* **16**: 1007–1015.

Zheng GX, Terry JM, Belgrader P, Ryvkin P, Bent ZW, Wilson R, Ziraldo SB, Wheeler TD, McDermott GP, Zhu J, et al. 2017. Massively parallel digital transcriptional profiling of single cells. *Nat. Commun.* **8**: 1–12.

Zhou Y, Jin R, and Hoi SCH 2010. Exclusive lasso for multi-task feature selection. In: *Proceedings of the thirteenth international conference on artificial intelligence and statistics.* JMLR Workshop and Conference Proceedings, pp. 988–995.
